# ActSort: An active-learning accelerated cell sorting algorithm for large-scale calcium imaging datasets

**DOI:** 10.1101/2024.08.21.609011

**Authors:** Yiqi Jiang, Hakki O. Akengin, Ji Zhou, Mehmet A. Aslihak, Yang Li, Oscar Hernandez, Sadegh Ebrahimi, Yanping Zhang, Hakan Inan, Omar Jaidar, Christopher Miranda, Fatih Dinc, Marta Blanco-Pozo, Mark J. Schnitzer

**Author notes:** Co-first authors, see the contribution statement at the end (Yiqi Jiang, Hakki O. Akengin, Ji Zhou). These authors co-supervised this work (Fatih Dinc, Marta Blanco-Pozo, Mark J. Schnitzer).

## Abstract

Recent advances in calcium imaging enable simultaneous recordings of up to a million neurons in behaving animals, producing datasets of unprecedented scales. Although individual neurons and their activity traces can be extracted from these videos with automated algorithms, the results often require human curation to remove false positives, a laborious process called *cell sorting*. To address this challenge, we introduce ActSort, an active-learning algorithm for sorting large-scale datasets that integrates features engineered by domain experts together with data formats with minimal memory requirements. By strategically bringing outlier cell candidates near the decision boundary up for annotation, ActSort reduces human labor to about 1–3% of cell candidates and improves curation accuracy by mitigating annotator bias. To facilitate the algorithm’s widespread adoption among experimental neuroscientists, we created a user-friendly software and conducted a first-of-its-kind benchmarking study involving about 160,000 annotations. Our tests validated ActSort’s performance across different experimental conditions and datasets from multiple animals. Overall, ActSort addresses a crucial bottleneck in processing large-scale calcium videos of neural activity and thereby facilitates systems neuroscience experiments at previously inaccessible scales.

## 1 Introduction

Systems neuroscience research has been revolutionized by large-scale Ca^2+^ imaging techniques, which now regularly produce terabyte scale datasets [1, 2, 3, 4, 5, 6, 7, 8, 9]. This rapid increase in neural data has necessitated automated methods for identifying neuronal sources, leading to the development of cell extraction algorithms [10, 11, 12, 13]. These algorithms facilitate large-scale analysis by detecting thousands of cells, sometimes up to a million neurons [9, 14, 15]. However, they can sometimes misidentify non-neuronal elements as neuronal structures, potentially leading to erroneous biological interpretations in downstream analyses [16, 17, 18].

To mitigate errors arising in automated cell extraction pipelines, experimental studies often perform quality control processes, known as *cell sorting*, to validate and refine cell extraction outputs [10, 11, 12, 13, 19]. In practice, researchers either laboriously annotate all cell candidates using specialized software (See CIATAH, [20]) or employ “cell classifiers” that are trained on a subset of annotated datasets to label the rest [10, 11, 20]. On the one hand, manual annotation is no longer practical given the large data sizes in modern neuroscience [8, 14, 15, 21, 22]. On the other hand, as we show in this work, existing automated methods still require substantial human labor to achieve desirable accuracy. Yet, human annotations are known to develop undesirable biases ([23, 24, 25, 26, 27]) due to, *e*.*g*., getting tired after long hours of repetitive work. Therefore, there is an urgent need for a scalable and robust solution that minimizes human labor without compromising the accuracy of the cell sorting.

A potential solution may lie in the active learning paradigm, a branch of machine learning that optimizes model performance while minimizing the need for annotated samples [28, 29, 30, 31, 32, 33, 34, 35]. In this paradigm, so-called query algorithms select the most informative and sparse samples to be labeled by human annotators [36, 37, 38]. To date, active learning has improved several scientific pipelines such as classifying pathological images [39, 40], detecting intracranial hemorrhage [41], and segmenting melanoma [42]. Inspired by these successes, we set out to leverage the active learning framework to accelerate the cell sorting process.

In this work, we introduce ActSort: an active learning accelerated cell sorting algorithm to process large-scale Ca^2+^ imaging datasets. As a first step, we developed a memory-efficient and user-friendly cell annotation software. Using this software, we designed an extensive cell annotation benchmark. This benchmark contains roughly 160,000 annotations of 40,000 cells collected using one- and two-photon mesoscopes across 5 mice, which we used to test and validate ActSort. Supported with novel features engineered by expert neuroscientists and a novel query algorithm tailored for cell sorting, ActSort matched human-level accuracies while reducing the required human labor to a negligible fraction of the full samples (down to 3% when training from scratch, and 1% if ActSort is initialized with previously annotated samples). In large-scale mesoscope movies, ActSort achieved convergence by sorting few hundreds of cells (c.a. 1 hour of human labor). In contrast, to achieve the same mark, labeling random subsets of representative datasets required sorting an order of magnitude larger number of cells. Notably, our analysis also revealed significant disagreements among human annotators, yet semi-automated classifiers mitigated this large variability across experimenters. Overall, our findings suggest that ActSort can save experimentalists significant time, and improve the reliability, robustness, standardization, and reproducibility of systems neuroscience research.

## 2 Results

Our cell sorting pipeline (ActSort) consists of three main components (Fig. 1): the preprocessing module, the graphical user interface (GUI), and the active learning module. The full pipeline is available to end-users with a point-click installation in our GitHub repository [43] and can be used as an application or in Matlab-online without a license. In this section, we introduce ActSort and validate it with several experiments on large-scale annotation benchmarks.

**Figure 1:**
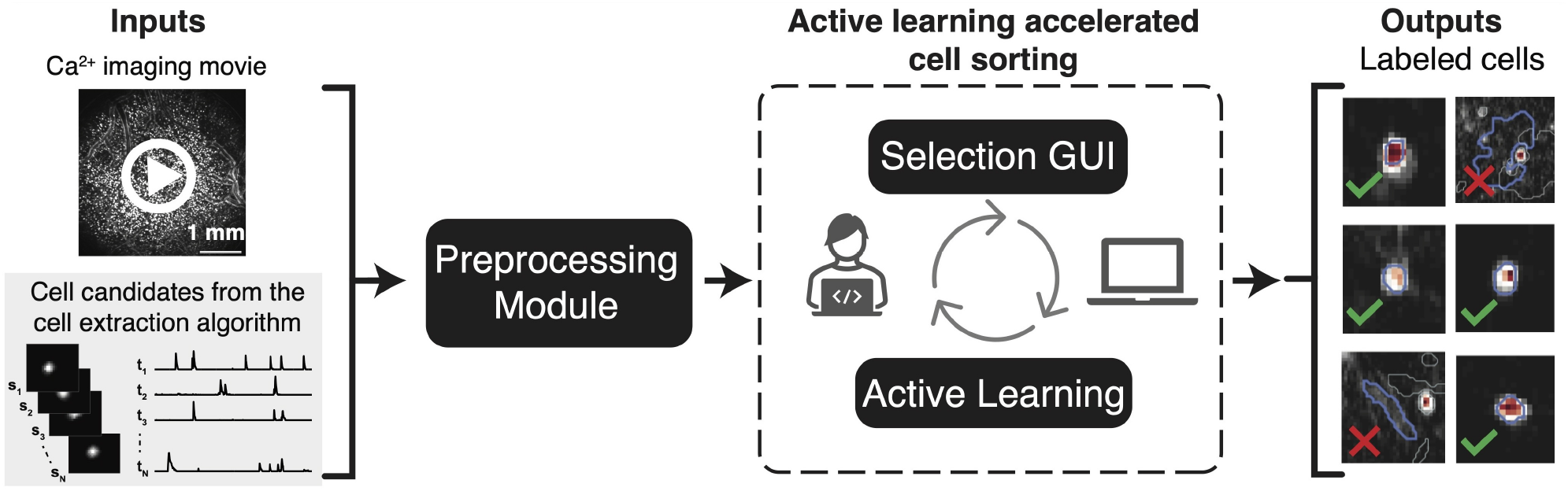
An active learning accelerated framework for rapid quality control of cell extraction results. We present ActSort, an active learning accelerated framework that requires minimal human labor to cull out false positive cell candidates extracted from large-scale neural recordings. *Left*. A typical preprocessing of Ca^2+^ imaging movies involves identification of cell candidates by automated cell extraction algorithms. ActSort is a cell sorting algorithm, which takes as an input the Ca^2+^ imaging movie and the cell extraction results from the prior steps, and culls out false-positives from the set of identified cell candidates. *Middle*. ActSort has three distinct modules. The preprocessing module jointly compresses the cell extraction results and the Ca^2+^ imaging movie to allow memory efficient cell sorting in large-scale Ca^2+^ imaging movies. The preprocessing module also computes quality metrics for each cell candidate (called “features,” **Methods**; Appendix A.1), which are later used in the active learning accelerated cell sorting module. The selection GUI allows human annotators to visually inspect the cell extraction results and annotate the true and false positive cell candidates (**Methods**; Appendix A.2). The active learning module, working in closed-loop feedback with the selection GUI, strategically selects next cell candidates to be annotated by the experimenter in order to optimally reduce the human effort. In the meantime, online cell classifiers are trained on the resulting annotated subset of the data. These classifiers are used as part of the active learning module and eventually automate the cell sorting process (**Methods**; Appendix A.3). *Right*. The resulting outputs from ActSort are probabilities and binary labels (cell or not a cell) for each cell candidate. Preferably, only small portion of the final outputs would need to be sorted by experimenters, whereas cell classifiers trained with the closed-loop active learning framework could confidently classify majority of the cell candidates.

### 2.1 Sorting cell candidates in massive datasets with a custom written software

Only five years ago, a Ca^2+^ imaging movie with a 100 GB size was considered large [12]. Today, neuroscientists routinely collect datasets several terabytes in size [44]. Given this rapid increase, we start our work by designing a scalable software with a primary focus on memory efficiency. Without keeping the full movie in RAM, the software needs to include essential information used by human annotators to determine whether a sample is a cell: i) the shapes of cells’ extracted spatial footprints, ii) their Ca^2+^ activity traces over time, and iii) the movie snapshots at the time of Ca^2+^ events (Fig. S1; Appendix A). While earlier work often retained only the first two elements for memory efficiency [10], we found that the spatiotemporal information provided by the Ca^2+^ movie snapshots was crucial for human annotators, especially in edge cases, which are particularly important for the active learning paradigm (see below). This design choice also aligns well with the newer Ca^2+^ imaging processing pipelines (see the cell sorting module in CIATAH [20]), and can be incorporated with an efficient data compression procedure without increasing dataset sizes significantly.

To achieve this design goal, as part of the preprocessing module (Fig. S2), we opted to compress the movie and the cell extraction results jointly. We saved the variables in sparse formats whenever applicable and retained only the relevant parts of the Ca^2+^ imaging movies where the cells had Ca^2+^ events (up to a pre-defined maximum; see Appendix A.1 for details). This approach allowed us to compress the large inputs of the traditional cell sorting pipelines (Fig. 1), resulting in approximately 270 ± 90 MB/1,000 cells (mean ± std over 5 mice) data sizes. Thanks to the extreme compression of the movie and cell extraction data, we were able to develop a memory efficient and user-friendly graphical user interface for cell sorting (Figs. S1, S2, and S3). This software does not require any coding knowledge, utilizes buttons and sliders to operate, and can be used on laptops.

Our software has an additional, originally unintended, benefit. Most public Ca^2+^ imaging datasets share cells’ spatial profiles and Ca^2+^ activity traces, but not the movies themselves due to large data sizes [45, 46]. Yet, motion artifacts and/or neuropil contamination can often not be identified by simply looking at Ca^2+^ activity traces, but can alter biological conclusions [13, 18]. Thanks to our memory-efficient data format, open-sourced datasets can now include relevant movie snapshots without astronomical increases in data sizes, and the quality of the Ca^2+^ activity traces can be probed with our software.

### 2.2 A cell sorting benchmark with 160,000 annotations across five mice

After coding the cell sorting software, we first searched for public datasets to validate the use of cell classifiers and active learning paradigm as a viable method for automated cell sorting. Although there are publicly available Ca^2+^ imaging movies with annotated cells, notably NeuroFinder challenge [47], these datasets are often region specific and contain few hundreds of cells. Yet, we have designed ActSort to process large-scale mesoscope movies. These modern movies simultaneously record large populations of neurons across multiple brain regions, and are now regularly collected by neuroscience labs [9, 14, 44]. To the best of our knowledge, no annotation dataset is publicly available with brain-wide mesoscope movies.

To bridge this gap, we curated a total of five mesoscope movies: i) three one-photon Ca^2+^ imaging movies spanning the half-hemisphere through a 7mm window (Table S1), ii) one Ca^2+^ imaging movie from previous work ([44]) spanning eight neocortical regions (Table S2), and iii) one two-photon Ca^2+^ imaging movie capturing layer 2/3 cortical pyramidal neurons in mice primary visual cortex (Table S3). We extracted an approximate total of 40,000 cells from these movies using the EXTRACT cell extraction algorithm [13]. We then had 6 annotators independently perform cell sorting on these movies, with 4 annotators per movie, totaling up to roughly 160,000 annotations (details in **Methods**; Appendix B). We used this benchmark to perform validation experiments in the rest of this work.

### 2.3 Automated cell sorting with linear classifiers utilizing newly engineered features

Automated cell sorting pipelines rely on training cell classifiers to annotate the unlabeled cell candidates [10, 20, 44]. These classifiers often use features derived from the cell extraction outputs and, in some cases, from Ca^2+^ imaging movies [20]. The central hypothesis behind this automation is that these derived features—such as the stereotypical shapes of cells’ spatial profiles—contain sufficient information to distinguish false positives from real cells. However, relying on a select few features overlooks a crucial aspect of cell sorting: human annotators incorporate information from broad sources in their decisions. Consequently, we hypothesized that cell classifiers would benefit from a broader class of features than those traditionally implemented in the literature.

To test this hypothesis, we first designed a traditional feature set (**Methods**; Appendix A.1), which aims to provide a strong baseline representing the set of features used by existing cell extraction pipelines [10, 12, 13, 20, 44]. With these features, we processed our half-hemisphere Ca^2+^ imaging movies from three mice, encompassing 28,010 cell candidates and 112,040 annotations (Table S1). Specifically, for each mouse, we trained cell classifiers using traditional features on half of the cell candidates and predicted the identity of the other half. These classifiers performed superbly in identifying true positives, correctly accepting 97% of the cells (data not shown). However, they were suboptimal in rejecting false positives, accurately rejecting only 67% of the candidates rejected by human annotators (Fig. 2**A**). Part of this low accuracy may be due to the fact that only a small fraction of the samples (often <10%) are false positives, and human annotators may struggle to remain consistent with their strategy for duplicate selection and/or when deciding edge cases. Nevertheless, this observation was in line with our prediction that there may be room for improvements in the feature sets used by cell classifiers.

**Figure 2:**
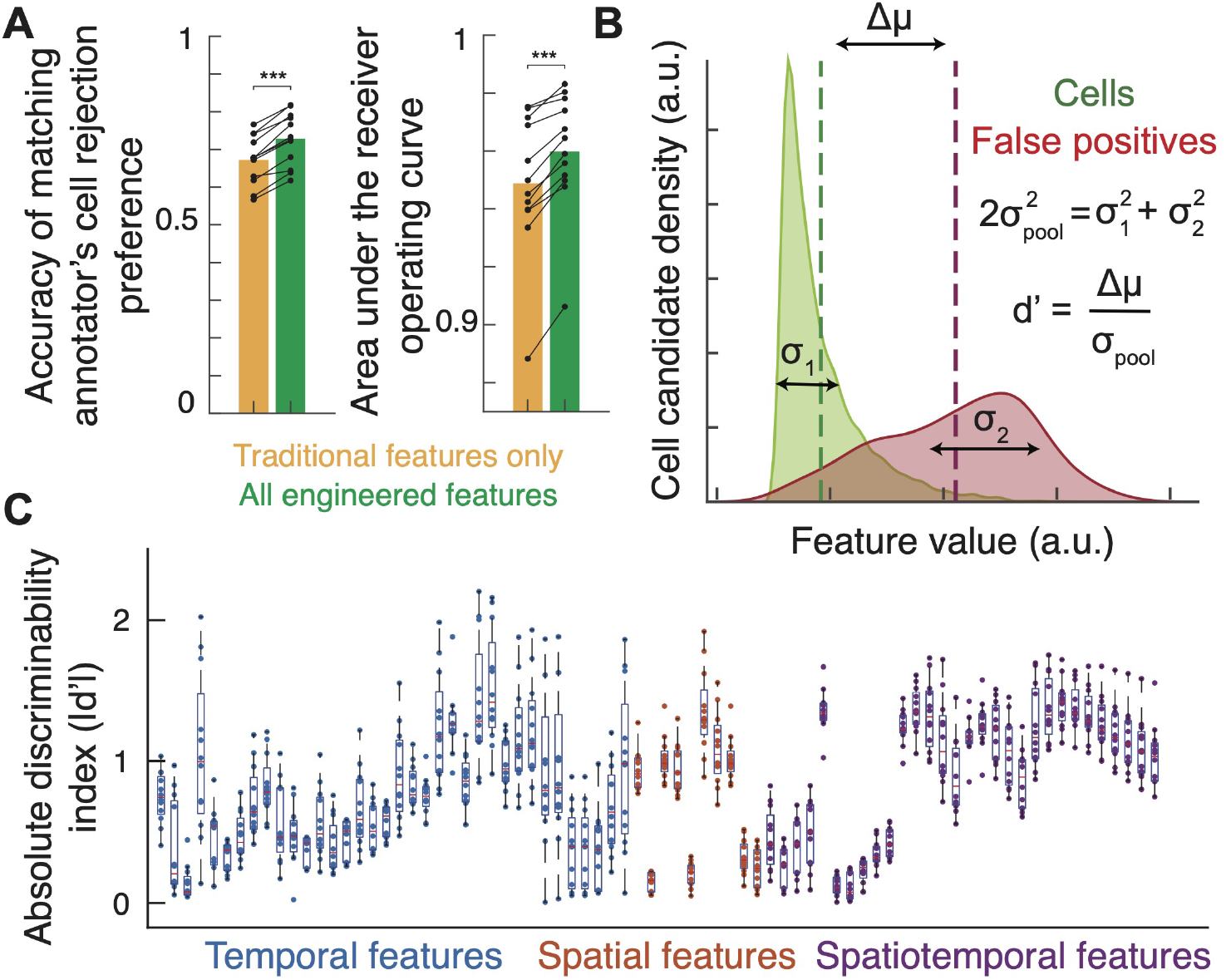
Feature engineering improves the accuracy of automated cell sorting with linear classifiers. To facilitate training of accurate cell classifiers (which label each cell candidate as *cell* or *not cell*), we engineered a total of 76 features. These features were computed from the compressed movie snapshots (Fig. 1), and cells’ spatial footprints and activity traces extracted from the Ca^2+^ movies (**Method**; Appendix A.1). For this figure, we used the half-hemisphere dataset with 3 mice and slightly more than 28, 000 cell candidates (Table S1). Each dot corresponds to an independent annotation instance. **A** We tested the cell classifier accuracies achieved with either the traditional or the full set of features. The former set refers to the simple features that are often used by extant cell extraction pipelines as quality metrics [10, 12, 13]. The newly engineered features increased classifiers’ ability to reject false-positives and led to improved area under the receiver operating characteristic curve, as quantified by two-sided Wilcoxon signed-rank tests (^***^*p* < 10^−3^). **B** To measure the effectiveness of individual features, we computed their discriminability indices (*d*′) that quantify how well each feature can separate out cells from false-positives. **C** Many features exhibited middle or strong effects (*d*′ ≥ 0.5 or *d*′≥0.8, respectively), whereas few features were not discriminative for this particular dataset. We confirmed that each feature had at least marginal contribution in at least one annotation instance across the full benchmark with diverse Ca^2+^ imaging movies (data not shown; multiple regression analyses, *p* < 0.05 for 74 out of 76 features, whereas *p* = 0.05 for the remaining two). Orange lines in box plots denote median values; boxes span the 25th to 75th percentiles. The whiskers extend to 1.5 times the inter-quartile range.

To bridge this gap, we engineered new features for the cell classifiers by mimicking the *spatiotemporal* information human annotators pay attention to during cell sorting. The full list of resulting 76 features can be found in Appendix A.1. This set includes traditional temporal features (e.g., number of spikes in a trace, signal-to-noise ratio) and spatial features (e.g., mean pixel intensity, circumference), as well as novel spatiotemporal features (e.g., severity of the non-Gaussian noise contamination). To ensure memory-efficient computation, these features were derived solely from the compressed data format, allowing feature extraction to be computed on demand or as part of the cell extraction routine. Re-analyzing the dataset from Fig. 2**A**, we found that the new features increased the rejection accuracy to 72% (Fig. 2**A**) without decreasing the accuracy of accepting true-positives (data not shown, 97%), leading to more effective separation between *cell* and *not cell* samples in our benchmarks (Fig. 2**B-C**).

### 2.4 Expert-engineered features set strong baselines against deep learning approaches

Deep learning approaches have seen recent success in relevant domains of our application [48]. In regards to cell sorting, a cell extraction pipeline, CAIMAN [12], trained deep classifiers to cull out false positives from two-photon calcium imaging datasets. These classifiers were trained solely on cells’ spatial footprint, not using any temporal or spatiotemporal information often used by human annotators. Consequently, this method was suboptimal compared to the consensus human annotation (see Table 1 in [12]), and cannot process one-photon imaging movies [49]. As we show below, neither is true for ActSort thanks to the generalization of features across broad imaging and experimental modalities. Yet, perhaps a better feature extraction may be performed in an unsupervised manner using deep learning approaches. Thus, as another strong baseline, we next investigated whether a pre-trained image model, a ResNet-50 architecture [50], could enhance the unsupervised discovery of features.

To perform this test, we extracted features from cropped cell images and movie frames using the pre-trained model (**Methods**; Appendix C.3). The movie frames, consisting of time-averaged snapshots around the cell’s location during Ca^2+^ events, offered spatiotemporal information, which was designed to provide a fair comparison to our engineered features. Using these images, we extracted 2,000 features from the penultimate layer of ResNet-50 [50, 51]. Notably, though extracted features led to increased true negative rate, on par with our engineered features, this came at the cost of significantly decreased true positives (down to ∼ 90%, Fig. S4). Therefore, due to their superiority (Figs. 2 and S4), we decided to use the newly engineered feature set in our ActSort pipeline.

### 2.5 Humans are inconsistent, but cell classifiers mitigate annotator biases

Can human annotations serve as ground truths for cell sorting? To answer this question, we next tested the possibility that human annotators may be inconsistent due to, *e*.*g*., personal biases. To quantify this inconsistency, we performed intraclass correlation (ICC) analyses [52], a method commonly used to assess the reliability of ratings in a scale (here, binary annotations), either by individual raters or averaged across multiple raters. Since, in experimental scenarios, only one human annotator typically handles a dataset, we computed a two-way mixed, single rater ICC [52], which measures the consistency of individual annotators rather than the group average. Across all five mice, the intraclass correlation scores for individual human annotators were *ICC* = 0.56 ± 0.06 (mean ± std), indicating that human annotators often disagree, and are inconsistent, on what they may classify as cells.

Next, we checked whether cell classifiers can mitigate individual biases and show higher consistency when trained on human annotations. To quantify the classifiers’ consistency, we split the annotation datasets into equal training and testing sets, trained the cell classifier on the former, and computed the label on the latter (See Appendix C.2). The cell classifiers had higher correlation scores, *ICC* = 0.64 ± 0.07 (mean ± std), showing improved consistency compared to individual human annotations *despite being trained on them*.

As the final test, we trained and evaluated the cell classifiers by reanalyzing a published one-photon Ca^2+^ imaging dataset covering the neocortex [44], which included 6,691 cell candidates and 26,764 annotations (Table S2). We once again computed the test probabilities from cell classifiers. These probabilities were compared with the empirical probability computed from annotator agreements (*e*.*g*., a vote of 2-2 corresponds to a 50% probability, Fig. S5). We found that annotator expertise was crucial for the accuracy of the cell classifiers trained with these annotations. But, as an overall trend, human annotators were overconfident in their choices. In contrast, the cell classifiers, especially those trained with more experienced annotators, were better calibrated (Fig. S5). Thus, we confirmed that cell classifiers with the newly engineered feature set can match or even surpass the consistency of individual human annotators.

Yet, if human annotators are inconsistent, how can we validate the accuracy of ActSort, or in general, automated cell sorting? To address this, we designed a cross-validated evaluation process that incorporated the fact that humans often disagree with each other (Fig. S6). Specifically, to evaluate the active learning routines trained with a specific annotator, we created a ground truth evaluator by taking the majority vote of the three held-out annotators. Fortunately, the majority votes among the held-out human annotators had good consistency with *ICC* = 0.79 ± 0.05 (mean ± std, two-way mixed, average measures [52]). Thus, throughout this work, the annotations performed by either the classifiers and/or the experimenters are evaluated on these majority-voted held-out labels.

### 2.6 Discriminative-confidence active learning query algorithm for cell sorting

So far, we introduced our custom software for cell sorting, discussed how to utilize feature engineering and cell classifiers to annotate unlabelled cell candidates, and established how to reliably and consistently evaluate the cell sorting results via majority votes. Now, we introduce the final piece of the puzzle: integrating the cell classifiers into the active learning framework with “query algorithms.”

In ActSort, the active learning query algorithm prompts human annotators to label specifically selected samples. These samples are expected to improve the automated cell sorting accuracy, which creates an online feedback loop between the annotator and classifier (Fig. 1). But how should the samples be chosen for annotation? A straightforward approach is to annotate a random subset and label the rest, as has been done in the literature to date [10, 20]. Yet, as we discuss below, better alternatives exist.

When designing alternative query algorithms, we focused primarily on speed and scalability, as sorting is performed on the spot and within seconds. Therefore, we considered query algorithms that can train and predict rapidly. Two such strategies are confidence-based active learning (CAL) [53, 54] and discriminative active learning (DAL) [55]. These query algorithms aim to mitigate two distinct types of uncertainties: (i) uncertainty regarding the sample’s position relative to the decision boundary of the current model (CAL), and (ii) uncertainty regarding whether labeled samples faithfully represent the full dataset (DAL). Both query algorithms achieve these aims with rapidly trainable linear classifiers (see **Methods** for details; Appendix A.3).

Briefly, CAL trains a cell classifier on the samples labeled up until that point and picks the unlabeled sample closest to the decision boundary for annotation. DAL trains a secondary “label” classifier to discriminate labeled cells from the unlabeled data and selects a sample most different from the already sampled subset. Evidently, an optimal solution should bring samples up for annotation that are close to the final decision boundary (CAL). However, first, the query algorithm should approximate where the final boundary should be with sparse samples. This, in turn, requires collecting diverse samples that represent, roughly, the full dataset (DAL). Thus, both approaches have their unique advantages.

To amplify the strengths of both approaches, we introduce a new query algorithm: discriminative-confidence active learning (DCAL). DCAL selects samples for human annotation by considering both the classifier’s uncertainty and the representation of samples in the unlabeled dataset. This is achieved by training both cell and label classifiers and adaptively combining their scores with a user-initialized weight *w* (**Methods**; Appendix A.3). As the number of samples increases, the weight is adjusted based on the accuracy of the label classifiers. Specifically, DCAL gradually approximates CAL, since the label classifiers tend to become random predictors due to the growing diversity of annotated samples. Meanwhile, the DAL component is crucial in the early stages to identify outlier data points (Fig. S7**A**). Thanks to the adaptive estimation process, DCAL depends minimally on the initial choice of the trade-off weight (Fig. S7**B**), and is thereby practically parameter free.

### 2.7 Geometrical interpretation of discriminative-confidence queries

The query algorithms we consider in this work – random, CAL, DAL, and DCAL – have simple geometric interpretations, which we discuss now using an example movie from our benchmark (Figs. 3 and S8). To illustrate the sample selection preferences by these query algorithms, we reduced the dimension of the feature space from 76 down to 2 using a partial-least squares analysis and plotted in Figs. 3**A** and S8.

**Figure 3:**
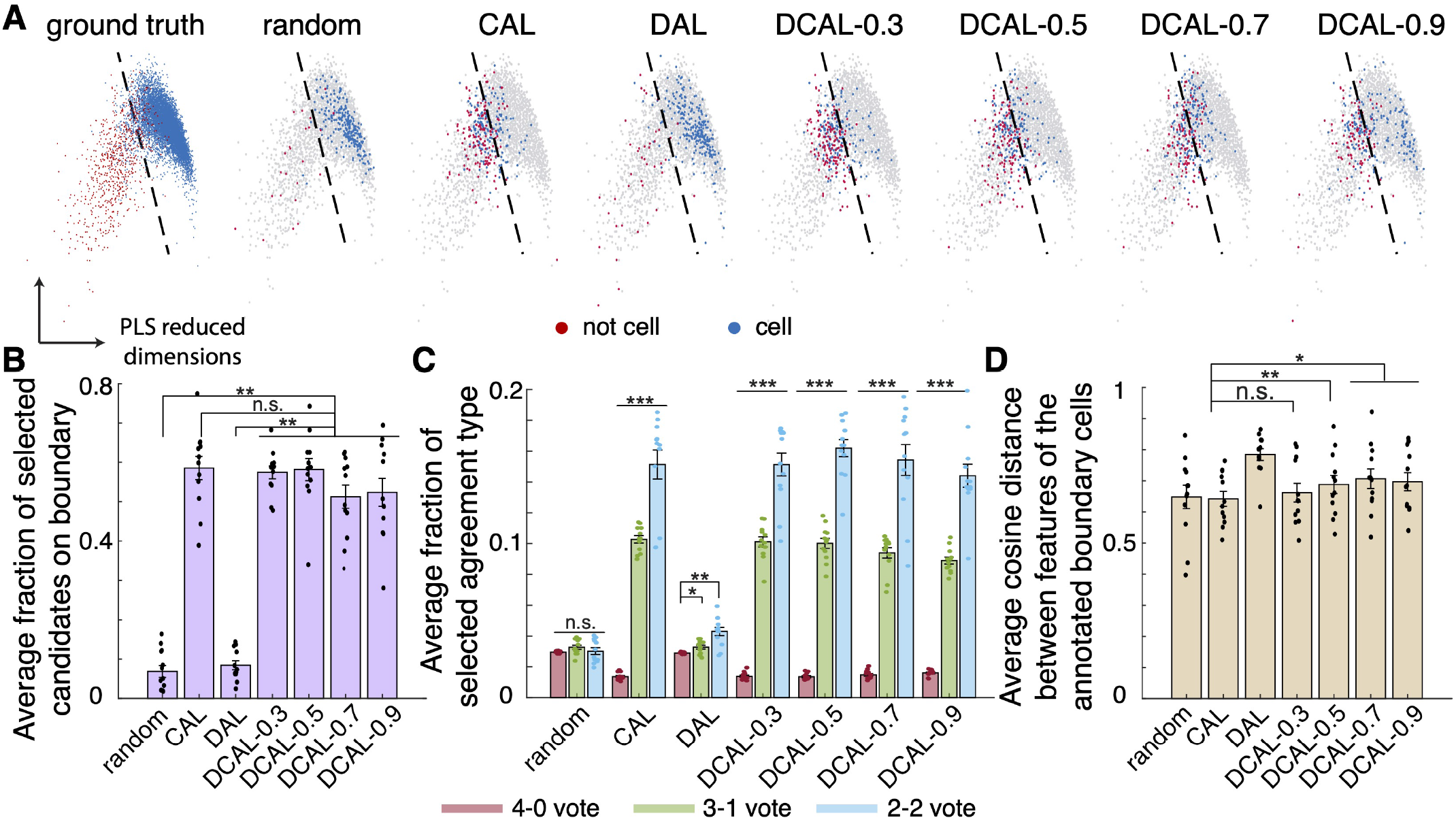
Discriminative-confidence active learning (DCAL) algorithm selects outlier boundary samples. Our active learning query algorithm (DCAL, **Method**; Appendix A.3) effectively addresses the limitations associated with the two prevalent query strategies, confidence-based active learning (CAL) and discriminative active learning (DAL). **A** Dimension reduction via supervised partial least squares (PLS) was applied to the 9,983 cell candidates from a representative annotation of the half-hemisphere dataset. For each query algorithm, we sorted up to 3% cell candidates and visualized them in the PLS-reduced dimensions computed from the majority-voted held-out labels. DCAL effectively identified the boundary cell candidates while simultaneously encouraging a broad coverage within the boundary. Blue dots represent cells (as annotated by a single human), whereas red dots are false positives. The dashed black lines correspond to (approximate) decision boundaries in the reduced feature space. The fraction next to DCAL indicates its user-defined initial weight. **B** The average percentage of the selected samples near the decision boundary (probabilities within [0.25, 0.75], **Method**; Appendix C.4) when sorting 3% of cell candidates with each algorithm. DCAL significantly outperformed DAL and random selection in identifying these candidates, and matched CAL, an algorithm designed solely to identify boundary samples. **C** The fraction of cell candidate types, based on annotators’ votes (0:4/4:0, 1:3/3:1, and 2:2), selected by each query algorithm normalized with respect to the total number in that particular confidence pool. DCAL and CAL prioritize the most ambiguous candidates for annotation process, which are particularly relevant for training linear classifiers as they lie on the decision boundary by definition. **D** The average cosine distance between standardized features of boundary samples, the same samples that lie within the decision boundary in **(B)**. DCAL enhances sample diversity, covering a broader feature space. In **(B-D)**, each dot represents a single annotation instance. Error bars: s.e.m. over 12 annotations. All tests are two-sided Wilcoxon signed-rank tests with Bonferroni-Holm corrections (^***^*p* < 10^−3, **^*p* < 10^−2, *^*p* < 0.05). See Tables S4 and S5 for additional details on the statistical analyses.

Both random and DAL queries comprised of diverse samples covering the full spectrum, with the latter showing increased diversity as expected. As a consequence, since there are for more true positive samples in the full dataset, both of these approaches picked mostly real cells up for annotation (Fig. 3**A** and S8; random and DAL). In contrast, CAL primarily picked cell candidates that are close to the decision boundary, leading to a more balanced selection of true and false positive annotations (Fig. 3**A** and S8; CAL). Yet, especially for sparse annotations, CAL had a preference to sample from higher density regions of the boundary (Fig. S8; CAL with 1%). DCAL, on the other hand, was practically equivalent to CAL in large annotation limit, but also mitigated the dense sampling with sparse annotation (Fig. 3**A** and S8; DCAL). In other words, DCAL picked *outlier* boundary cells, providing a comprehensive coverage across the boundary.

Next, we quantified these observations by focusing on a subset of the dataset, which we termed “boundary samples” (**Methods**; Appendix C.4). Here, we confirmed that both CAL and DCAL selected a high percentage of boundary samples (Fig. 3**A, B**). These selections preferentially included samples that were challenging for human annotators to agree on (Fig. 3**C**). In selecting these samples, DCAL has shown significant improvements over DAL (Fig. 3**B,C**), which was expected since the latter has no relevant incentive. Notably, however, DCAL also matched the performance of CAL in picking boundary samples, an algorithm solely designed to do so, and had improved diversity in selected boundary cells (Fig. 3**A, D**). Thereby, these results jointly confirmed that DCAL preferentially picked samples lying on the decision boundary and those further from previously picked samples, *leading to broader coverage and better representation of the boundary* (Figs. 3 and S8).

### 2.8 Performance evaluations of ActSort on the human annotated benchmarks

Though automated cell sorting is a necessity, work to date has not performed a clear validation of such approaches in large scale benchmarks. Similarly, though active learning query algorithms may provide improvements, existing approaches to date have focused on simple random sampling strategies. To perform the first test of this paradigm, we ran several experiments on our benchmarks using the cross-validated evaluation process (Fig. S6). Specifically, we tested the cell classifiers and the active learning query algorithms (random, CAL, DAL, and DCAL) with closed-loop simulated experiments, which mimicked the real-time annotation that ActSort aims to facilitate (Fig. 4**A**).

**Figure 4:**
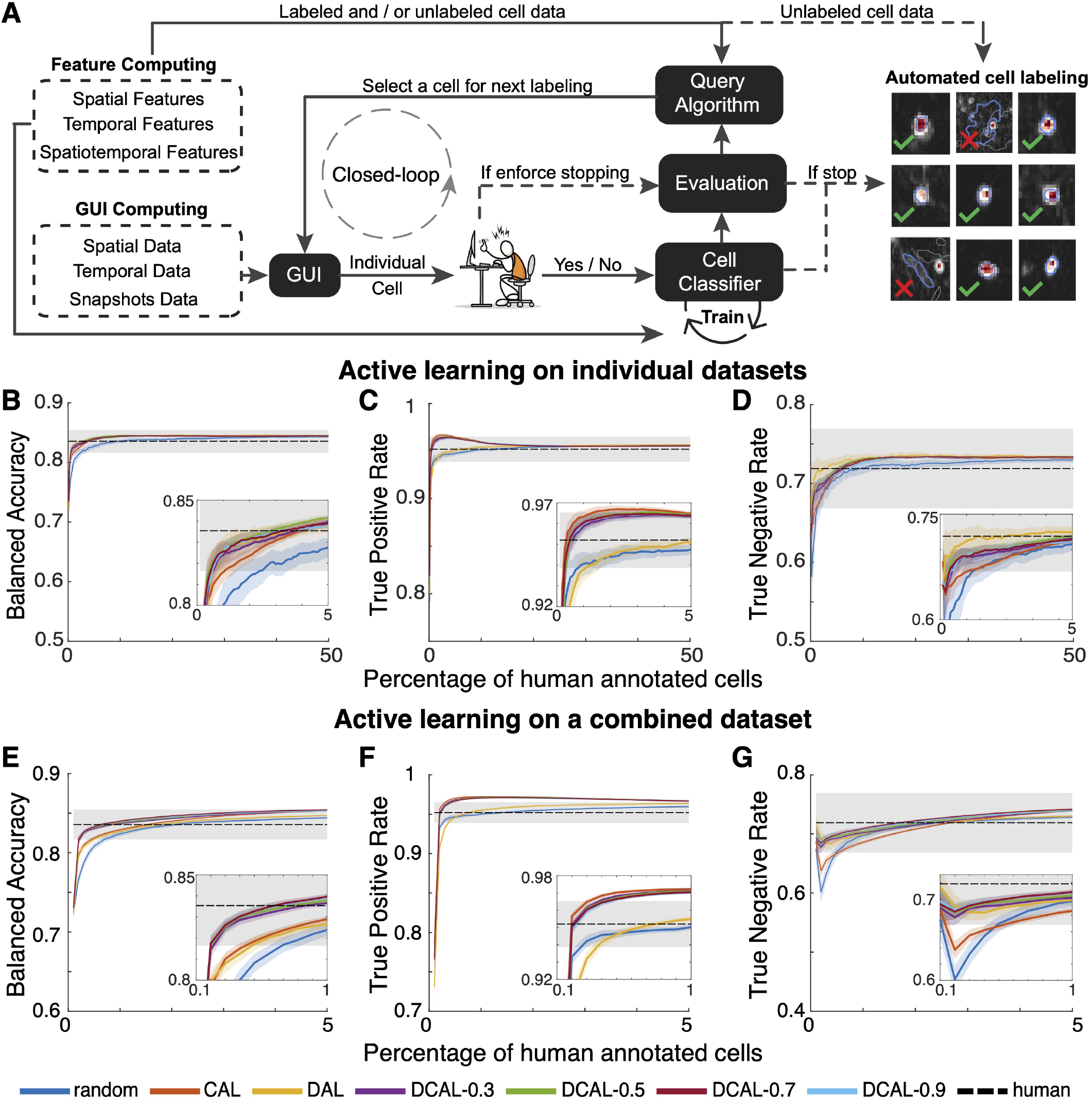
ActSort converges rapidly (at 5 − 10% data) and outperforms human annotators (at 1 − 3%). To assess the efficiency and effectiveness of active learning accelerated cell sorting, we simulated online annotation scenarios using the half-hemisphere datasets from three mice and the associated 12 human annotations (Table S1). **A** ActSort creates a feedback loop between the experimenter and the query algorithm. Human annotator uses a graphical user interface (GUI) and labels a cell candidate selected by the query algorithm as either *cell* or *not cell*. Online cell and label classifiers (**Methods**; Appendix A.3) are then trained on the labeled samples, number of which grows with each iteration. Using these classifiers, the query algorithm selects new candidates for human annotation, thereby closing the feedback loop. These steps iterate until sufficient amount of samples is annotated. Then, manual sorting ceases and the remaining cell candidates are automatically labelled using the predictions of the trained cell classifier (**Methods**; Algorithm S1). **B-D** Using the half-hemisphere dataset (Table S1), we ran active learning experiments individually on each annotation instance (out of 12 annotations across three mice), sorting up to 50% of cell candidates. We evaluated various query strategies - random, CAL, DAL, and DCAL — by computing **(B)** balanced accuracies, **(C)** true positive rates, and **(D)** true negative rates as a function of the annotation percentages. CAL and DAL either preferred high true positive or true negative rates, respectively, at the expense of the other. In contrast, DCAL achieved the desirable trade-off. Notably, with only 3% annotated data, DCAL matched the human annotator performances. In comparison, random sampling required up to 10% annotation to achieve the same mark. Solid lines: means. Shaded areas: s.e.m. over 12 annotators. Dashed line and gray area: mean and s.e.m. from 12 human annotations. **E-G** We re-ran the experiments in **(B-D)**, but for a scenario where multiple imaging datasets are processed simultaneously. To achieve this, we combined annotations across the three Ca^2+^ imaging movies by taking the combinations of 4 annotations per movie. This led to a total of 64 augmented annotators, each labelling c.a. 28,000 cell candidates, and corresponding ground truths from held-out majority votes. In this case, both CAL and DAL were suboptimal and had almost similar accuracies to the random sampling, presumably due to the complexities introduced by combining cell candidates from different mice. In contrast, DCAL outperformed all other strategies and reached human-level balanced accuracies with < 1% of the annotated samples. Solid lines: means. Shaded areas: s.e.m. over 64 annotators. Dashed line and gray area: mean and s.e.m. from 64 augmented human annotations. In all cases, cell classifiers had lower variability compared to human annotations, despite being trained on them, as indicated by the smaller shaded areas. For additional metrics computed in these experiments, see Tables S6, S7, and Fig. S13.

As a first test, we processed individual Ca^2+^ imaging movies covering half-hemisphere from three distinct mice (Table S1 and Fig. 4**B-D**). With as little as 5% annotations, DCAL achieved human level true negative rates (Fig. 4**D**), whereas all algorithms except DAL and random sampling had an above human level true positive rate of ≥ 95% with ≤ 1% annotated samples (Fig. 4**C**). As expected, DCAL’s performance was mostly independent from the chosen weight due to the adaptive estimation process (Appendix A.3), but also distinct (and better) compared to either DAL or CAL. Notably, ActSort with DCAL reached human-level balanced accuracy with annotating roughly 3% of the full dataset and converged within < 10% beyond these levels, *even though the classifiers were trained on human outputs* (Fig. 4**B**). In comparison, random sampling reached the same marks with roughly 10% and 50% annotated samples, respectively (Fig. 4**B-D**).

Next, we considered the scenario where cell candidates from multiple animals are processed in a single combined batch (Fig. 4**E-G**). To test this scenario, we augmented 64 new annotators and corresponding evaluators by selecting one annotator from each dataset (Fig. S6**B**). Consequently, each combined dataset contained slightly more than 28,000 samples across three mice. In this experiment, with < 1% annotated instances, DCAL reached human-level balanced accuracy, and had minimal improvements beyond roughly 3% annotated samples (Fig. 4**E**). Surprisingly, both DAL and CAL massively underperformed DCAL, offering little improvement over random sampling (Fig. 4**E-G**). These results underscore the importance of broader coverage and better representation of the decision space, especially when the annotation dataset contains diverse cell candidates sampled from multiple imaging sessions.

The half-hemisphere-wide movies we considered so far can be considered as high quality cell extraction results, with false-positive rates already as low as 5-10% per movie and high quality imaging and experimental conditions. Next, we asked whether ActSort would be able to accurately process lower quality imaging datasets. To test this, we processed one of the cortex-wide movies from [44] with less stringent quality parameters in EXTRACT such that the resulting cell extraction output had many false-positives, around 35%, by design (Fig. S9). We also tested ActSort on a two-photon Ca^2+^ imaging movie with residual motion (Fig. S10). In both datasets, DCAL outperformed all other approaches. Hence, the low false positive rate in the cell candidates or the optimal imaging/processing conditions cannot explain the success of ActSort in converging rapidly with few annotated samples. We further confirmed that these results were robust to changes in hyperparameters, such as the regularization parameter (Fig. S11) and classifier thresholds (Fig. S12).

### 2.9 ActSort with pre-labeled samples generalizes across animals

As a test of ActSort’s generalization ability, we conducted additional experiments with a set of pre-labeled samples using the three Ca^2+^ imaging movies of the half-hemisphere dataset. Specifically, we pre-trained the cell classifier on one dataset using 50% of the pre-labeled samples. This classifier was later fine-tuned using the real-time annotations on a new dataset from a different mouse. To test this pre-labeling approach, we paired 3 movies with each other, creating six dataset pairs. Each pair had 16 augmented annotations, pre-labels came from one of four annotators of the first movie, fine-tuning was performed using one of four annotators of the second movie (Fig. 5**A**). The details of the fine-tuning process can be found in Algorithm S2 and Appendix C.5.

**Figure 5:**
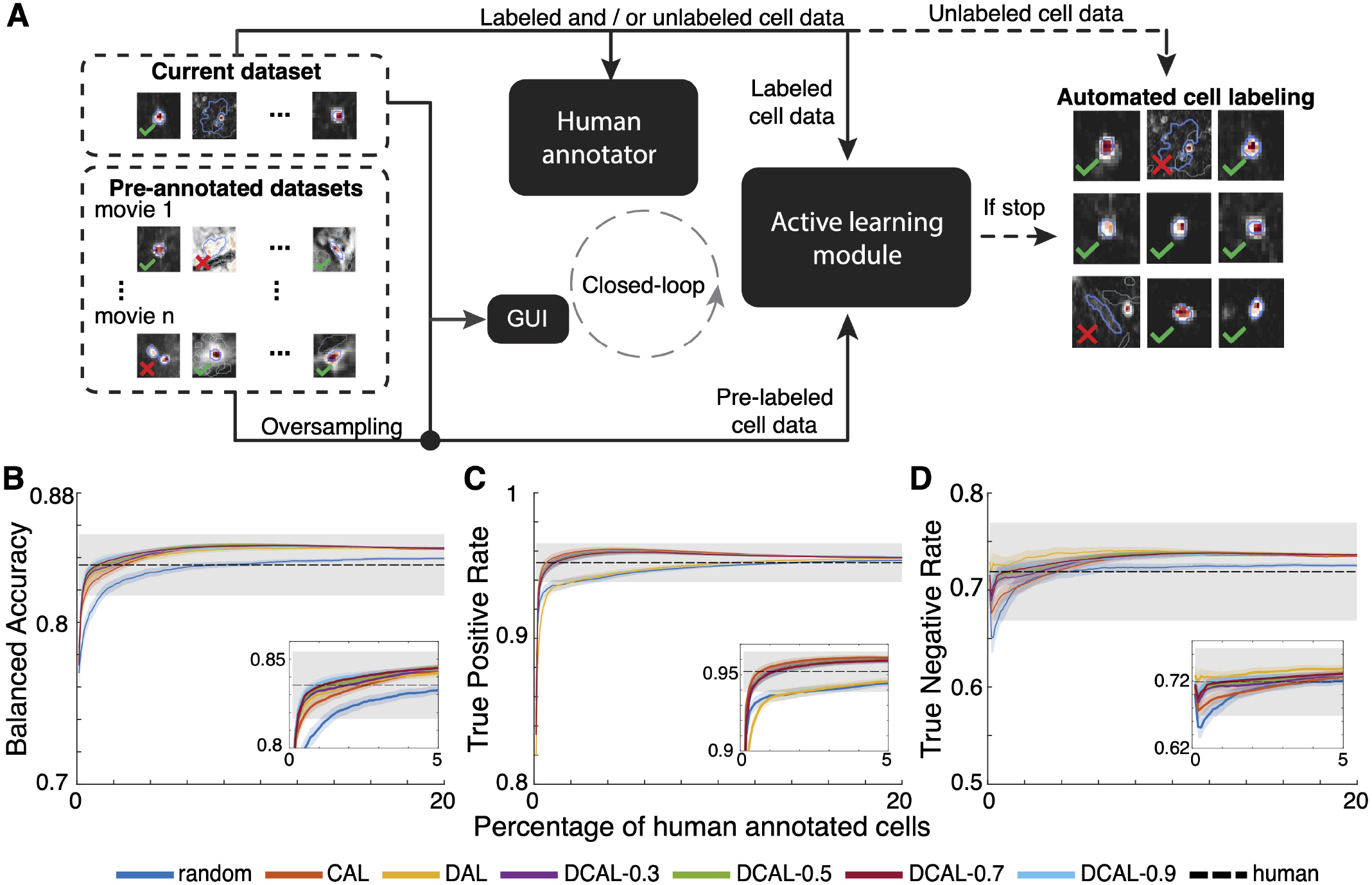
ActSort pre-trained with previously annotated datasets converges faster on new animals. We tested ActSort’s generalization abilities by pre-training classifiers on data from already annotated mice. These classifiers can be fine-tuned to new datasets, accelerating the convergence and decreasing the required human labor. **A** To test ActSort’s generalization capacity, we once again used the half-hemisphere datasets from our benchmark. Specifically, we trained cell classifiers on half of the cell candidates from one mouse (representing pre-annotated datasets) and fine-tuned on samples from another mouse (representing the new dataset that needs to be annotated). In this fine-tuning scenario, the GUI displays the information on the current dataset, whereas the active learning module trains the cell classifiers using both newly annotated and pre-annotated data in the background. The query algorithm then selects the next candidates for human annotation, facilitating fine-tuning to the dataset of interest (**Method**; Appendix C.5). **B** - **D** We created 6 pairs of mice, involving 16 augmented annotations per pair, totaling 56,020 cell candidates and 224,080 annotations. The first dataset is sorted up to 50% of cell candidates. Using the same query algorithm, the second dataset is sorted up to 20%. We evaluated various query strategies - random, CAL, DAL, and DCAL - by computing **(B)** balanced accuracies, **(C)** true positive rates, and **(D)** true negative rates by averaging over the 96 data-annotation pairs. DCAL achieved human level performance with as little as 1% annotation, three times faster compared to training without pre-labels (Fig. 4**B-D**). Solid lines: means. Shaded areas: s.e.m. over 96 augmented annotations pairs. Dashed line and gray area: mean and s.e.m. from human annotations. See Table S8 and Fig. S13 for additional details.

Our results, illustrated in Fig. 5, demonstrate the effectiveness of combining pre-labeled samples with DCAL. Notably, with pre-labels, DCAL achieved a human-level balanced accuracy with as little as 1% annotated samples and converged with roughly < 6% annotations (Fig. 5**B**). In contrast, random sampling did not achieve human-level performance until c.a. 8% annotations and had not converged even with the 20% samples (Fig. 5**B**). Similar to before, we found that DAL had similar performance to random sampling in picking true positives (Fig. 5**C**), for which CAL was the better algorithm. However, DAL outperformed CAL in the true negative rates (Fig. 5**D**). As their combination, DCAL often outperformed or matched the accuracies of both of these approaches, highlighting its robustness and effectiveness in sorting cells under varying conditions (Fig. 5).

### 2.10 ActSort culls out non-neuronal signals that may correlate with behavior

Taken together, our benchmarking experiments demonstrate the critical role of active learning query algorithms in automated cell sorting. DCAL, which optimally balances two canonical active learning approaches, consistently outperformed human annotators, sometimes with as few as < 1% of annotated samples. Yet, a final question remains: Is cell sorting truly necessary?

To address this, we re-analyzed previously published recordings from the dorsomedial striatum of freely behaving mice [56]. Each dataset includes neural calcium activity recorded using the fluorescent calcium indicator GCaMP6m in spiny projection neurons of either the direct (9 mice) or indirect pathway (10 mice) of the basal ganglia (dSPNs and iSPNs, respectively). These datasets, originally processed using an independent component analysis-based cell extraction algorithm (ICA, [57]), contained 336 ± 132 (mean ± std) cell candidates, which were sorted by human annotators. We re-sorted these datasets using ActSort, with human annotations as inputs and CAL as the query algorithm due to the low diversity in cell candidates. Due to the dataset’s nature, the evaluation was performed against a single annotator, yielding a balanced accuracy of 90 ± 4% (mean ± std) for ActSort annotations, converging at 20% annotations.

As shown in Fig. 6, both human and ActSort-sorted dSPNs and iSPNs significantly predicted animal speeds. Remarkably, ActSort achieved the same level of predictive accuracy with as little as 1% annotation, resulting in cells with nearly identical speed predictions (Fig. 6**B-C**). However, even the rejected cell candidates contained residual information about the animals’ speeds, likely due to brain motion artifacts. This highlights a crucial issue: non-neuronal signals in calcium imaging movies can inadvertently correlate with animal behavior, leading to potentially misleading biological conclusions. Thus, the false-positives should be culled out to prevent unsolicited information leakage in the downstream analyses.

**Figure 6:**
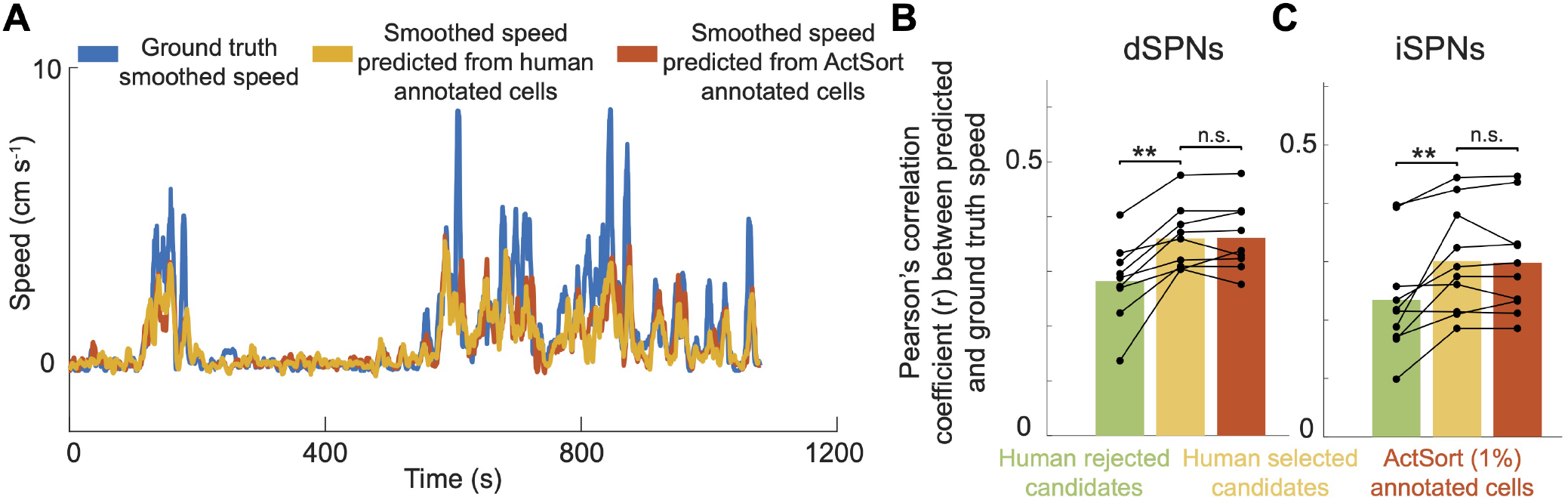
Culling false-positives is necessary to prevent spurious Ca^2+^ signals leaking into data analysis. As a test of whether omitting cell sorting can lead to spurious biological conclusions, we studied Ca^2+^ imaging data that was previously acquired in the dorsomedial striatum of freely behaving mice using a head-mounted, epifluorescence miniature microscope [56]. To test whether the human annotated and the ActSort predicted cells (trained with CAL and using only 1% annotation for each mouse separately) provided similar levels of speed decoding, we trained support vector machines to predict the (smoothed) speeds of the animals. **A** Visualization of speed decoding from an example animal. **B-C** In both dSPNs **(B)** and iSPNs **(C)**, after controlling for cell numbers, human annotated cells provided better speed decoding compared to human rejected cells, and were equivalent to ActSort sorted cells. More importantly, however, cells that were rejected by human annotators were informative regarding the speed of the animal. This means that in brain regions not coding for speed, cell extraction artifacts, likely due to brain motion, can encode animal’s behavior spuriously, potentially leading to incorrect biological conclusions. In **B, C**, Each dot corresponds to results from a single animal, averaged across 1,000 speed decoders trained from distinct train-test splits of the data. All tests are two-sided Wilcoxon signed-rank tests (^***^*p* < 10^−3, **^*p* < 10^−2, *^*p* < 0.05).

## 3 Discussion

ActSort is as a standalone quality control software, which can be used to probe the outputs of *cell extraction pipelines* such as EXTRACT [13], Suite2p [10], ICA [57], CAIMAN [12], and others [11, 19, 58, 59]. Such existing cell extraction algorithms take the raw Ca^2+^ movie as their inputs, correct the brain motion, perform simple transformations to standardize the Ca^2+^ imaging movies, identify putative cells’ spatial profiles and temporal activities, and often perform primitive quality controls to output the final set of neural activities. Historically, additional quality controls on the cell extraction outputs would be performed with manual annotation, which was feasible for the small neural recordings with up to thousands of neurons [23, 24, 25, 26, 27, 20]. Yet, with the advent of large scale Ca^2+^ imaging techniques, now recording up to one million cells [9], manual review is no longer realistic. Instead, the field of experimental neuroscience direly needs automated quality control mechanisms that would correctly identify the true cell candidates while rejecting false positives misidentified by the extraction algorithms.

ActSort is the first scalable and generalizable solution in this direction. Yet, previous works (as parts of existing cell extraction pipelines [10, 12, 13]) had tackled this problem in specific instances: Suite2p designed cell classifiers based on some basic features to increase the precision of the algorithm [10], CAIMAN pre-trained a deep classifier for two-photon movies [12] (though not applicable to 1p Ca^2+^ imaging movies [49]), whereas EXTRACT performed thresholding on a set of quality metrics [13]. Notably, these existing automated methods with pre-trained cell classifiers often found success only for high quality 2p Ca^2+^ imaging movies [10, 12], and even then underperformed human annotators [12]. One-photon Ca^2+^ imaging datasets, on the other hand, are quite diverse in their imaging (miniscope vs mesoscope) and experimental (head-fixed vs freely behaving) conditions and face additional challenges due to low resolution and high background noise. With ActSort, we sought a generalizable solution that does not target a specific modality or require substantial re-training, but uses interpretable features that are robust across various behavioral experiments, Ca^2+^ indicators, imaging conditions, and techniques.

To provide a baseline for existing methods in our benchmarks, we designed a feature set called “traditional features” (Fig. 2), including features used by classifiers in these prior works (See Appendix A.1). Moreover, these methods did not use active learning, instead annotated only randoms subsets. Thus, in our experiments, these prior methods (or a plausible upper bound to be more exact) are represented as the random sampling query algorithm, which uses the full feature set to allow fair comparisons to active learning approaches. In all benchmarking experiments, we observed the clear advantages of using active learning query algorithms (specifically, DCAL) as opposed to random sampling, as is traditionally done in the literature [20, 44]. The most striking observation, however, is the significantly lower variability of classifier predictions compared to human annotations despite being trained on them (see, *e*.*g*., the differences between gray and colored shaded regions in Figs. 4 and 5). Thus, our results suggest that the automation is expected to not only reduce the human labor, but also standardize it.

Though we performed extensive analysis to highlight the efficiency and effectiveness of ActSort, our work is merely a first step for what may hopefully become a fruitful collaborative subfield comprising of experimental neuroscientists and active learning researchers. There are several future directions that future work could improve upon, which we will briefly summarize below.

Firstly, in this work, we used linear classifiers for rapid online training during cell sorting and reproducibility of annotation results. This was mainly rooted in the fact that pre-training deep networks required substantial data and standardization across various Ca^2+^ imaging movies. We believe that with additional public datasets that may follow our lead, this direction can become reality (as was the case for a different, yet relevant, problem of spike extraction [60]). Our results in this work have set a strong baseline for such future deep-learning approaches.

One limitation of ActSort is that it comes as a quality control pipeline to existing cell extraction approaches. Therefore, any mistakes in the cell extraction step would automatically propagate to cell sorting. Yet, some of these mistakes can be mitigated or highlighted post hoc. For instance, future ActSort versions could involve options for merging segmented or duplicated cells, or identifying motion in activity traces and thereby notifying the user to improve their preprocessing steps.

Another important aspect requiring further exploration is the expertise level of the annotators. To date, each Ca^2+^ imaging movie is often annotated by a single annotator, who unfortunately can tire after long hours and become inconsistent as we have discussed above. This was the reason behind our choice to have the movies in our benchmark be annotated by multiple researchers. Yet, the human annotators had different levels of expertise in working with Ca^2+^ imaging datasets. Hence, a potential moderation relationship can exist depending on the expertise of the annotator. To fully explore this relationship requires further research with *more* annotators per dataset, by having the same annotators sort the same cells in different times, and/or testing sorting effectiveness before and after a standardized sorting training.

Finally, in this work, we validated ActSort for one-photon and two-photon Ca^2+^ imaging datasets and for sorting binary classes. Yet, our framework is generally applicable to other modalities, barring additional feature engineering performed by domain experts in those fields, or with a pre-trained deep network. For instance, by replacing the logistic regression with a multinomial version and approximating CAL scores with entropy instead of decoder scores, our work readily applies to multi-class datasets that may jointly include, *e*.*g*., dendritic and somatic activities. With correct features, the framework we introduced here should also be helpful for the newly emerging voltage imaging recordings, especially as the number of cells will inevitably increase in such movies with the technological advances.

Overall, while human annotation was possible in datasets discussed in this work due to the manageable number of cell candidates, data analysis and hypothesis testing at scale with millions of neurons will require robust and automated cell sorting tools. In such cases, ActSort, and tools that will follow its lead, are essential for filtering out non-neuronal artifacts that could otherwise compromise the accuracy and reproducibility of neuroscience research.

## 4 Conclusion

In this work, we introduced ActSort, a quality control algorithm for processing large scale Ca^2+^ imaging datasets. We utilized real-time human feedback via a closed-loop active learning paradigm to decrease the need for humman annotation by two-orders of magnitude. We achieved this by first engineering features that generalize across experimental and imaging conditions, and by developing a simple, yet effective, active learning query algorithm. Finally, to support the development of active learning algorithms in systems neuroscience, we introduced the first publicly available benchmark for cell extraction quality control on both one- and two-photon large-scale Ca^2+^ imaging, comprising five datasets: four one-photon and one two-photon Ca^2+^ imaging datasets, with approximately 40,000 cells and 160,000 annotations.

The potential impact of this work is two-fold: i) vastly decrease the number of hours humans spend annotating modern Ca^2+^ imaging data sets and ii) surpass the accuracy of cell classification and biases by experts. A paradigm requiring that merely 1-5% of cells be annotated by a human can reduce cell sorting from several hours per Ca^2+^ imaging data set, to merely tens of minutes. Through ActSort, researchers can reclaim tens of hours that would have otherwise been spent labeling modern massive data sets. Importantly, increasing cell detection accuracy can be largely beneficial to modern systems neuroscience, which relies on minimizing false positive cell identification to avoid spurious results. Thus, removing the human element from repetitive tasks that can be prone to errors can ultimately improve the reliability, robustness, and reproducibility of systems neuroscience research.

## Acknowledgements

We would like to thank Beyzanur Arican Dinc for suggesting the intraclass correlation analysis to quantify the consistency among annotations. HOA acknowledges funding support from Ozyegin University. JZ thanks Matlab’s Toolbox Trainee program for fostering a connecting to the EXTRACT team at Schnitzerlab. CM acknowledges funding support from the Stanford University Wu Tsai Neurosciences Institute Interdisciplinary Scholar Award. FD, YL, and YJ received funding from Stanford University’s Mind, Brain, Computation and Technology program, which is supported by the Stanford Wu Tsai Neuroscience Institute. MJS gratefully acknowledges funding from the Howards Hughes Medical Institute, NIH Brain Initiative, Simons Collaboration on the Global Brain, and the Vannevar Bush Faculty Fellowship Program of the U.S. Department of Defense.

## Data and code availability

ActSort software and the benchmark used in this study are publicly at our Github repository: https://github.com/schnitzer-lab/ActSort-public.

## Methods

### A Software details

In this work, we introduce ActSort, a user-friendly pipeline consisting of 3 modules: a preprocessing module, a cell selection module, and an active learning module. *The preprocessing module* efficiently reduces the data size associated with 10, 000 neurons down to few GBs, which includes not only cells’ Ca^2+^ activity traces, but also spatial profiles, and movie snapshots during Ca^2+^ events -allowing end users to check the quality of the data-, together with the engineered features. This compact representation allows the ActSort pipeline to be run locally in laptops, despite original movie sizes of up to few hundred GBs. The joint compressing of the movie snapshots and cell extraction information offers another key contribution to the neuroscience community with implications for sharing imaging datasets publicly. *The cell selection module* features a custom design with an easy-to-use interface that displays temporal, spatial, and spatiotemporal footprints, and incorporates a closed-loop online cell classifier training system. During the annotation process, the software provides real-time feedback by displaying predicted cell probabilities and the progress made by the human annotator as well as the fraction of unlabelled cells that ActSort is confident about. This allows experimentalists to monitor their work and make informed decisions regarding automated cell sorting outputs. *The active learning module* works in the background, strategically selecting candidates for human annotations, and trains cell classifiers with annotated candidates.

#### A.1 Preprocessing module

The preprocessing module is included within the GUI. This module reduces repetitive calculations by precomputing all necessary components.

The inputs for the module are the Ca^2+^ imaging movie and the corresponding cell extraction output, which includes spatial data for each cell in the dataset and trace activity data. This module performs 1) a joint compression of Ca^2+^ imaging movies and cell extraction results, 2) computes all the visualization components, and 3) performs feature engineering. The output for the module is a ‘sorting file’ that contains the necessary components for cell visualization in GUI, features for each cell, and an ‘info’ field that provides useful statistics.

To accommodate different system capabilities, the user can select whether data is loaded entirely into memory or is read in chunks. Furthermore, for large datasets, parallel processing on both CPU and GPU can be selected. Note that for small datasets, parallel processing might slow down this process.

##### A.1.1 Joint compression of Ca^2+^ imaging movies and cell extraction results

For efficient handling of the growing sizes of cellular data on local laptops (with limited memory capabilities), we designed a new method for compressing both imaging movies and cell extraction results.

###### Compression of Ca^2+^ Imaging Movies

Traditionally, cell sorter GUIs require the Ca^2+^ imaging movie to be loaded along with cell extraction results. However, the growing size of these movies makes it impractical to store and load them both on hard disk and RAM. To mitigate this, we implemented Ca^2+^ event detection and snapshot capturing steps in our precomputation process.

1. **Ca**^2+^ **Event Detection**: Using the trace activity we obtain from the cell extraction algorithm, we identify the five largest Ca^2+^ events for each cell that represent distinct high activity moments. We utilize a publicly available spike detection algorithm (peakseek.m [61]) to determine these Ca^2+^ events.
2. **Snapshot Capturing**: For each selected Ca^2+^ event, a cropped snapshot is extracted comprising ten frames in a small window that includes neighboring cells during that time. Only these movie snapshots are stored. This process eliminates the need to load or share the entire movie file.

###### Compression of Cell Extraction Results

ActSort requires two main components from the cell extraction process: spatial filters and trace activities of each cell.

The spatial filters are typically stored in a three-dimensional array ([Height x Width x Number of Cells]), where each layer contains the spatial filter of one cell. Given the small size of each cell, this array is sparse and requires significant memory to store and load into RAM. To address this, we convert the spatial filters array into a sparse matrix, which allows to save a great amount of memory. All future code is organized to ensure this sparse matrix is never converted back to its full form and computations are performed directly on the sparse matrix. Additionally, trace activities are converted from ‘double’ to ‘single’ data type to save memory but still keep the precision at the same time.

##### A.1.2 Visualization Components Calculation

On top of the Ca^2+^ imaging movie snapshots and the compressed spatiotemporal information from the cell extraction algorithm, we also compute several components for visualization purposes in GUI:

- **Cell Centers**: Stores the center coordinates of each cell.
- **Ca**^2+^ **Event Indices**: Indices of the five most informative Ca^2+^ events for each cell’s trace activity.
- **Cell Boundaries**: Contains the boundary coordinates of each cell in spatial data.
- **Snapshots**: Stores snapshot images taken from the movie.
- **Snapshot Filters**: Filters applied to the snapshots.
- **Snapshot Traces**: Traces corresponding to the snapshots, used for feature extraction.
- **Snapshot Cell Boundaries**: Boundary coordinates of cells in the snapshots.
- **Snapshot Contrast Limits**: Proper contrast limits to better visualize the snapshots.
- **Neighbor Boundaries**: Boundaries of neighboring cells in the snapshots.
- **Maximum Projection**: The maximum intensity projection of the movie, used as a background image during cell label visualization.

##### A.1.3 Feature engineering

Below is the list of features we used for this work. The traditional ones are colored in blue.

> *Features 1 to 36 are based on the Ca*^2+^ *trace activities (activity signals of the neurons detected by the cell extraction algorithm) of the given cell dataset*.

Feature 1 calculates the signal-to-noise ratio for the given trace activity. A lower signal-to-noise ratio indicates a noisy data, which is more likely to contain false positives. A higher ratio indicates a higher quality data. This feature is taken from EXTRACT [13].

Feature 2 calculates how much a cell’s trace activity signal differs from its smoothed version. If the difference is high after smoothing, then it is likely for the cell trace activity to contain a good amount of noise. This feature is taken from EXTRACT [13].

Feature 3 calculates the number of weak Ca^2+^ events within the trace. We categorize the Ca^2+^ events as follows: We compute the 90th quantiles of the Ca^2+^ activity for each cell, and divide them by half. Any Ca^2+^ event with lower Ca^2+^ amplitude is considered “weak,” whereas the ones with higher Ca^2+^ amplitude are considered “strong Ca^2+^ events.”

Feature 4 calculates the standard deviation of the upper top 10% of each Ca^2+^ activity trace, revealing the variability within the Ca^2+^ event amplitudes.

Feature 5 measures the average time interval between two consecutive strong Ca^2+^ events. Constant and rapid spiking is often associated with motion artifacts and/or blood vessels.

Feature 6 measures the average time interval between two consecutive weak Ca^2+^ events. Feature 7 computes the average time interval between two consecutive Ca^2+^ events.

Feature 8 measures the average length of all Ca^2+^ events in a trace activity. Stronger Ca^2+^ events are a good sign of good candidates.

Feature 9 measures the average width of Ca^2+^ events. Bad candidates tend to have Ca^2+^ events that last longer.

Feature 10 calculates the standard deviation of the Ca^2+^ event width within a trace of cell activity. A more widely distributed Ca^2+^ event width can be a good indicator of noisy and less precise activity.

Feature 11 calculates the standard deviation of the Ca^2+^ event width among the strong Ca^2+^ events. This can serve as an indicator of their quality. A narrower distribution of Ca^2+^ event widths suggests a higher level of consistency.

Feature 12 measures the proportion of total activity in a cell trace made up of significant Ca^2+^ events rather than noise, referred to as relative power. This metric helps the distinction between actual Ca^2+^ events (consistent signal energy) and the likelihood of noise.

Feature 13 calculates the proportion of cell trace activity that surpasses the noise threshold. This tells the frequency of cell activities that is distinguished from the background noise.

Feature 14 counts the instances when the cell’s activity surpasses the noise threshold significantly. This provides insights into the frequency of firing or notable cell activities.

Features 15 - 29 are derived from different scales and threshold values: 3, 10, 30, and 100, respectively— for the calculation of relative power (Feature 12), the proportion of the cell trace activity that surpasses the noise level (Feature 13), and the number of strong Ca^2+^ events (Feature 14).

Features 30 - 34 calculate a quality score that indicates how much of a neuron’s activity occurs within certain frequency ranges of the spectrum. A higher score indicates that the majority of the neuron’s activity is happening within the specified frequency range, suggesting that the data quality is good and that the neuron is reliably detected. The frequency ranges are determined by lower and higher threshold parameters. In our work, the thresholds are set to define four specific ranges within the frequency spectrum: the lowest 5% (0-0.05), the range from the lowest 5% up to the highest 95% (0.05-0.95), the lower half (0-0.5), and the upper half (0.5-1).

Feature 35 calculates for each cell the proportion of other cells in the dataset with which it has a high correlation, using a similarity threshold of 70%. This essentially measures how similar a cell’s activity is to the activity of the other cells in the dataset.

Feature 36 finds the total fluorescence that was caused by the highest 10% activity in a given cell activity trace. This metric can be useful in detecting unusual neuron activity that doesn’t reflect a common neuron behavior in the dataset.

> *Features 37 to 46 are based on the spatial filters (the overall pictures of the cells according to the cell extraction algorithm) in the given cell dataset*.

Feature 37 calculates the total surface area of each cell in the movie. This can be useful in detecting cells that are too large or too small in size.

Feature 38 finds the number of distinct bodies that a cell candidate has. Cells that are detected to have more than one body tend to be false positives.

Feature 39 calculates the total sum of pixel values within each cell’s spatial filter, which is a measure of the total activity or brightness. ‘Bad’ cells tend to have unusually high or low activity.

Feature 40 measures the circumference of the cell. Typically, cells with too large or too small circumference tend to be false positives.

Feature 41 measures the distance from the cell center to the nearest edge of the movie’s field of view. Cells that are located closer to the edges tend to be noise or artifacts due to disruptions in the movie, such as boundary effects or uneven illumination.

Feature 42 computes the circularity of a given cell, which measures how closely its shape aligns with a perfect circle. Cells with a higher degree of circularity are generally preferable candidates. This feature is taken from EXTRACT [13].

Feature 43 computes the eccentricity of a given cell, which indicates how stretched out a cell’s shape is. Candidates with higher eccentricity values tend to be ‘bad’ cells. This feature is taken from EXTRACT [13].

Feature 44 computes the average pixel value of the given cell, which is a measure of the average activity. ‘Bad’ cells tend to have unusually high or low activity.

Feature 45 calculates the spatial corruption, which is known as errors or inconsistencies within the spatial filter of the cell by evaluating the local variance relative to the overall variance of the cell image. This feature is taken from EXTRACT [13].

Feature 46 calculates the maximum spatial correlation of each cell with all other cells in the movie. This is helpful to identify cells that have similar activity patterns. This feature is taken from EXTRACT [13].

> *Features 47 to 76 are based on spatial filters and trace activities of the cells that are captured by the cell extraction algorithm, along with key movie frames that correspond to the highest points of cell activity. We call these spatiotemporal features*.

Features 47 - 50 evaluate the alignment between the cell’s spatial filter, obtained from the cell detection algorithm, and critical movie frames extracted from the actual movie during the cell’s highest activity time intervals. Feature 47 averages the errors to identify cells with pronounced deviations. Feature 48 seeks the minimum error to check the accuracy of the closest match. Feature 49 uses the 10th percentile to detect rare but significant errors. Feature 50 uses the 20th percentile for a broader to avoid missing detections in the alignment of spatial filters and key movie frames.

Features 51 - 66 calculate the correlations of activity within the neuropil area -a dense network of neurons and their components that can cause false positives when activated-by analyzing the interactions between neighboring pixels in and around the cells. This feature is taken from EXTRACT [13].

Features 67 - 76 calculates the correlation between the cell activity traces and movie frames. This feature is taken from EXTRACT [13], but is not used as a quality metric within EXTRACT pipeline.

#### A.2 Graphical user interface (GUI)

We developed a user-friendly graphical user interface to visualize the cell data. We added many useful tools to assist users in annotating cells. The memory optimization performed by the preprocessing module allows to use of this GUI in practically any local computer.

Below is the list of functionalities within the ActSort GUI, which we illustrate in Fig. S3:

– **Cell Navigation**: Includes slider bars, buttons, and a number field to navigate through cells within the dataset.
– **Cell Stats**: General statistics show the distribution of cells categorized as good or bad, as well as the number of cells that have been labeled and those that remain unlabeled.
– **Zooming Options**: Includes a zoom slider and a zoom cursor for scanning and zooming into specific areas.
– **Contrast Sliders**: There are sliders to adjust the minimum and maximum contrast values for both the main map and snapshots.
– **Cell Color Transparency**: An option that allows the user to decide an appropriate contrast for the cells’ spatial profiles.
– **Cell Border Color Options**: Depending on the dataset, providing an option to change the cell border color can help users better visualize the cells on the main map.
– **Hiding Cells**: An option to hide sorted cells, both good and bad, which is helpful for visualization especially when working with large datasets.
– **Click to Select**: An option to select cells by clicking on them in the map.
– **Multiple Selection**: An option that allows selecting multiple cells from the map.
– **Sorting Unsorted Cells Only**: An option, when selected, already sorted cells are no longer shown to the user when moving back and forth with the slider.
– **Classifier Threshold**: An option to adjust the classifier threshold for prediction.
– **Predict Cells Button**: An option that predicts the identities of the cells based on the decision thresholds (see below)
– **Model selection**: The user can change any of the active learning algorithms, CAL, DAL, or DCAL.
– **ActSort Decisions**: Whether ActSort labels the current annotation as a cell or not.
– **ActSort Statistics**: The probability of the current annotation being a cell, assigned by ActSort.

#### A.3 Active learning approaches

We define a binary *cell classifier, h*_*θ*_, between *cell* and *not cell*, with input data 𝒳 ∈ ℝ^*N* × *K*^ and binary ground truth labels 𝒴 ∈ ℝ^*N*^, in which *N* is the total number of extracted putative neurons and *K* = 76 is the dimension of the features. Second, we defined a *label classifier, p*_*ϕ*_, between labeled data, ℒ ^(*t*)^ = {(*x*_*i*_, *y*_*i*_)} for 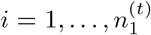, and the unlabeled data, 𝒰^(*t*)^ = {*x*_*i*_} for 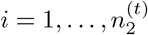, at a given iteration *t* such that 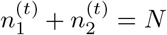, forming a new dataset 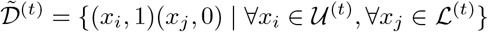. The former classifier is responsible for both the final outputs and the query algorithm, whereas the latter is used by the query algorithm only. To allow real-time training, and owing to its already sufficient accuracy thanks to the new feature set (Fig. 2**A**, also see below), we used logistic regression for both classifiers:

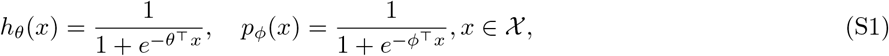

which were trained using the loss function (same for both classifiers)

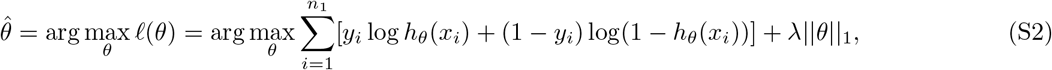

where *λ* is the L1 regularization strength^1^. The overall online closed-loop active learning pipeline is explained in Algorithm S1.

In this work, we tested four strategies as query algorithms, random sampling, confidence-based active learning (CAL) [53, 54]; discriminative active learning (DAL) [55]; the newly developed discriminative-confidence based active learning (DCAL). The overall training structure is depicted in Algorithm S1.

##### Algorithm S1 ActSort

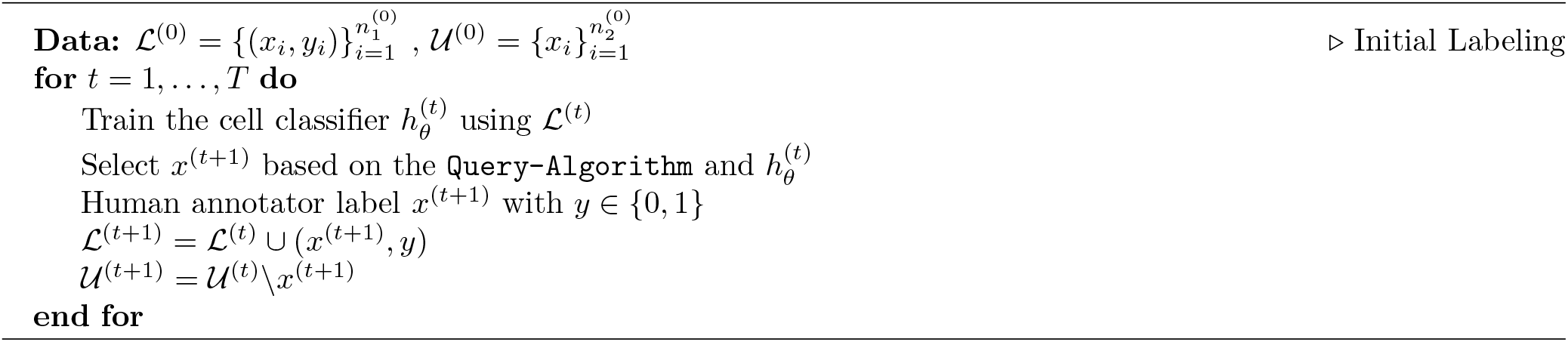

##### Traditional baselines

The random sampling algorithm randomly selects an unlabeled sample *x*^(*t*)^ from 𝒰. After human annotation, this sample is moved to the labeled set ℒ ^(*t*+1)^ and the cell classifier is updated accordingly:

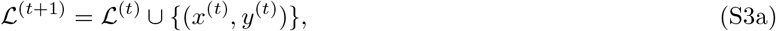

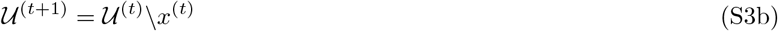

##### Confidence-based active learning

The CAL algorithm selects a sample according to the uncertainty sampling decision rule, which identifies unlabeled items that are near a decision boundary in the current cell classifier model. In a multi-class setting, one can use entropy to define the uncertainty of the sample points in the unlabeled dataset, which is how we shall motivate the CAL algorithm. Let ℋ (·) represent the entropy of a set of samples. In round *t* + 1, we want to choose the sample that produces the maximum reduction in entropy given the trained cell classifier 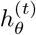 in round *t*:

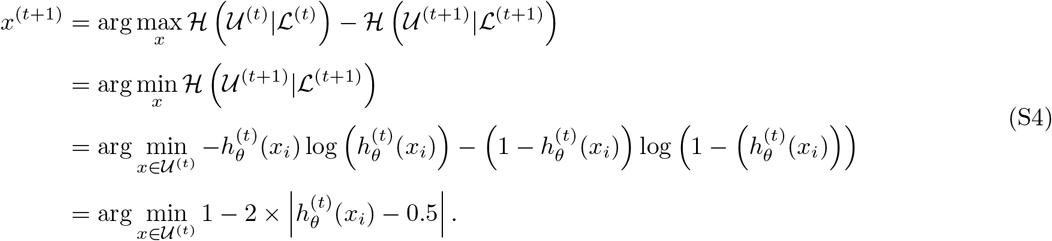

In this work, since we are using only two classes, the probability distribution that maximizes the entropy is the one where *h*_*θ*_ is as close to 0.5 as possible. However, extension of ActSort to multi-class problems, for example one that includes the identification of dendrites, could benefit from the more general entropy minimization framework.

##### Discriminative active learning

The DAL algorithm considers a different type of uncertainty, *i*.*e*., how well a data point is represented in the labeled pool [55]. Here, one constructs a new label space, 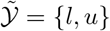, that specifies whether the data sample is from the labeled set or the unlabeled set. For example, in round *t*, the training dataset becomes 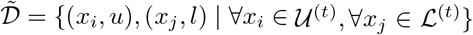. For each round of the active learning process, we train a binary classifier,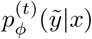, to predict whether the samples in 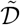 are labeled or not. Then, one selects the sample from the unlabeled set that satisfies

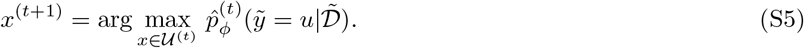

In this work, we modified the original DAL algorithm from [55] to improve its performance on our benchmark by training the classifier 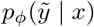 with random oversampling of the minority class samples in the dataset 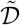.

##### Discriminative-confidence active learning

In order to achieve a query score for the DCAL algorithm, we define an uncertainty-score, 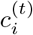, and a discriminative-score, 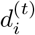, corresponding to the two types of uncertainty, for each iteration *t* as:

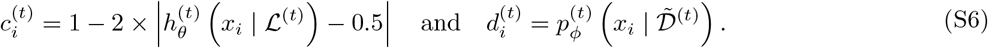

Empirically, once the labeled set accurately represents the full data, annotating candidates at the decision boundary yields the most significant improvements (See Fig. 3). Thus, we select the sample that maximizes the adaptively weighted sum of the uncertainty and discriminative scores from the unlabeled dataset:

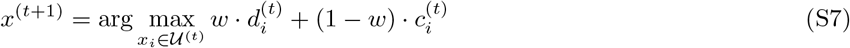

where *w* is an adaptive weight defined as

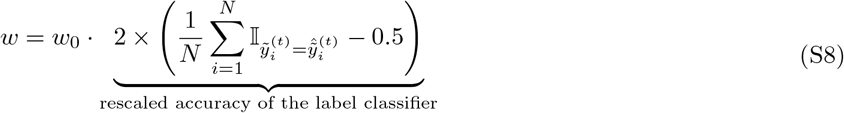

where *w*_0_ is the pre-defined weight, 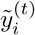 is the ground truth label and 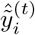 is the predicted label of the label classifier for the sample *i*. This adaptive estimation of *w*, in which we multiply the original weight *w*_0_ with the rescaled accuracy of the label classifier, allows DCAL to revert to CAL once the discriminative classifier is just providing random labels. This leads to convergence to the same performance between CAL and DCAL for larger *t*, and preferentially picks outlier *boundary* cells, ignoring outlier cells that are not relevant for the classification. In this work, we refer to DCAL algorithm with the initial pre-defined weight *w*_0_ as DCAL-*w*_0_.

### B Benchmark details

We provide five large-scale Ca^2+^ imaging datasets, totaling 40,000 cells with 160,000 annotations. These datasets include three one-photon Ca^2+^ movies spanning the half-hemisphere through a 7mm window (Table S1), one one-photon Ca^2+^ movie spanning eight neocortical regions (Table S2), and one two-photon Ca^2+^ movie focused on layer 2/3 cortical pyramidal neurons (Table S3).

For the half-hemisphere dataset, we used triple transgenic *GCaMP6f-tTA-dCre* mice from Allen Institute (Rasgrf2-2A-dCre/CaMK2a-tTA/ Ai93). To prepare mice for in vivo imaging sessions, we performed surgeries while mice were mounted in a stereotaxic frame under anaesthesia. We created a cranial window by removing a 7*mm* diameter skull flap over the right cortical area S1 and surrounding cortical tissue. We covered the exposed cortical surface with a 7*mm* diameter glass coverslip. A custom wide-field fluorescence macroscope with a field of view covering the full cranial window was used for neural activities imaging. For epifluorescence illumination, we used a LED with spectrum centered 475nm. We acquired Ca2+ videos of neural activity (50 Hz frame rate, 1, 708 × 1, 708 pixels) on the fluorescence macroscope. We used EXTRACT to extract the neurons from these movies [13].

To test ActSort on datasets with inflated false-positive rates, we used previously published imaging datasets imaging eight neocortical brain regions [44]. We performed cell extraction with weak quality checking [62, 13] on one of the sessions of roughly 20 min duration, which led to 2345 false-positive regions of interest.

To record the two-photon Ca^2+^ imaging movie, we employed a custom two-photon mesoscope with a multi-foci illumination technique [63]. This system facilitated recordings over an extensive field-of-view (2 × 2 *mm*^2^) at a frame rate of 30 Hz and a resolution of 2 *µ*m pixels (1024 × 1024 pixels). Utilizing this equipment, we captured a movie of layer 2/3 cortical pyramidal neurons from the primary visual cortex in live triple transgenic mice from the Allen Institute. The movie was extracted using the EXTRACT pipeline [13].

With this benchmark, we tested five different active learning algorithms: random, CAL, DAL, and our newly developed algorithm, DCAL. We compared the performance of these algorithms on the three types of datasets using the following methods: 1) the canonical active learning workflow (Fig. 4**A-D**), 2) batch processing (Fig. 4 **E-G**), and 3) cross-mice fine-tuning (Algorithm S2, Fig. 5). We evaluated performance using seven metrics: balanced accuracy, true positive rate, true negative rate, area under the receiver operating characteristic curve, recall, precision, and F-score, as a function of the percentage of human-annotated samples. The results for the canonical active learning using the half-hemisphere dataset are shown in Fig. 4**B-D** and Fig. S13**B** (Table S1). Batch processing results for the half-hemisphere dataset are presented in Fig. S4**E-G** and Fig. S13**C**, while the cross-mice fine-tuning results are depicted in Fig. 5 and Fig. S13**D**. The results for canonical active learning using the neocortical brain regions dataset (Table S2) and the two-photon microscope dataset (Table S3) are shown in Fig. S9 and Fig. S10, respectively.

Moreover, for the three experiments on the half-hemisphere dataset, we also provided tables summarizing the experiments’ performance for the canonical active learning (Table S6), batch processing (Table S7), and cross-mice fine-tuning experiments (Table S8).

### C Empirical experiments details

The experiments for large-scale Ca^2+^ imaging cell sorting were conducted on an Intel Core i9 processor, utilizing parallel for loops in MATLAB. These experiments took 5 days to sort 50% of the cells across 3 datasets, each involving 4 annotators, with 10 repetitions. For batch processing experiments, we used the same Intel Core i9 processor, running sequentially in MATLAB. This process required 2 days to sort 3% of cells from a total of 28,000 samples, with 6 repetitions. The cross-mice fine-tuning experiments were executed on Stanford Sherlock, utilizing MATLAB. These experiments were run in parallel across different methods and took 3 days to complete, with 3 repetitions. All cell classifier is a logistic regression with *h*_*θ*_(*x*) ≥ 0.65 predicted as *cell*, and *h*_*θ*_ < 0.65 predicted as *not cell*, see Appendix C.4 for the justification.

#### C.1 Estimating cell candidates’ labels with cell classifier on full dataset

For a dataset containing *N* cell candidates, denoted as 𝒟 = {*x*^(1)^, *x*^(2)^, …, *x*^(*N*)^} and each cell candidate has feature dimension *d, i*.*e*., *x*^(*i*)^ ∈ ℝ ^*d*^. To estimate the predicted categorical probability for each cell candidate by the cell classifier *h*_*θ*_, we conducted an alternate train-test split. Firstly, we split the dataset into set *A* and set *B*, each with 50% amount of data. Secondly, we train the cell classifier *h*_*θ*_ on dataset *A* and estimate the categorical probability for each cell candidate 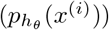 in set B. Next, we alternate set *A* and set *B* by training with data from set *B* and predict the label for each cell candidate in set A. Combining the second and third steps, we obtain the estimation of the categorical probability for each cell candidate without information leakage between the training and testing datasets. The final estimated categorical probability for each cell is by averaging over the 10 repetitions.

#### C.2 Picking the cell classifier thresholds for classification

Human annotators are prone to reject candidates that lie on the decision boundary during the annotation process. This tendency is due to the highly imbalanced nature of the dataset and the annotators’ preference for avoiding artifacts, even if it means missing a few actual cells. Therefore, using the default threshold of 0.5 does not accurately reflect the actual cell sorting process.

To find a threshold that aligns with human annotation criteria, we tested various threshold values ranging from 0.1 to 0.95 as shown in Fig. S12 using the three half-hemisphere datasets. The cell classifier labels a candidate *x* as *cell* if *h*_*θ*_(*x*) ≥ *β*, and as *no cell* if *h*_*θ*_(*x*) < *β*, where *β* is a predefined threshold that can be selected by human annotators. A smaller *β* results in a higher true positive rate, indicating more cells are correctly identified. On the other hand, a higher *β* results in a higher true negative rate, indicating more artifacts are correctly excluded.

From Fig. S12 **B, E**, and **H** we observe that, with 5% annotated samples, *β* < 0.8 is best for correctly accepting all cells, while *β* > 0.65 is best for correctly removing all artifacts. The precision graph (Fig S12**K**) shows that *β* = 0.65 most closely matches the human annotators’ inherent threshold. Thus, we set the classifier threshold *β* to 0.65 for all our experiments to best reflect the human annotation process.

#### C.3 Feature Extraction Experiments with ResNet-50 Model

Deep learning has shown great potential in representation learning across various fields. However, training a deep neural network is time-consuming and heavily dependent on the size and diversity of the training data, as well as the standardization across the data. Brain recordings, particularly 1-photon (1p) imaging videos, present significant challenges due to their diversity in background, imaging and experimental conditions, and cell shapes/types/sizes. In this experiment, we compared our expert-engineered features with those derived from a deep-learning approach. We used ResNet-50 for feature extraction and classification on 1p calcium imaging snapshots. Specifically, we provided the extracted cell profiles and the average cropped movie snapshot, resized to 224 × 224 pixels and converted from grayscale to RGB, averaged over frames with cellular activity, to ResNet-50. We then collected 2,000 features from the final layer.

Our engineered features outperformed the deep learning approach, which, although surprisingly effective at rejecting cells (and outperforming traditional features), did not surpass our engineered features as shown in Fig. S4. The fact that deep learning-based feature extraction was less successful on 1p movies aligns with conclusions from prior research [12, 60]. Additionally, even this relatively simple experiment (extracting features from average movie frames with ResNet-50) took approximately 30 minutes for 10,000 cells using an NVIDIA RTX 4090 GPU, and around 2 hours without one, which exceeds our design constraints.

We contend that our expert-engineered features, combined with the provided public dataset, establish a valuable benchmark for the cell quality control process.

#### C.4 Definition of boundary samples

The boundary samples for a logistic regression classifier are defined as the samples that have predicted probability between 0.25 and 0.75: {(*x, y*) | 0.25 < *h*_*θ*_(*x*) < 0.75}. The predicted probability for each sample is estimated by an average of the test set’s predicted probability from 10 repeats. The test set’s predicted probability is obtained by i) splitting the dataset equally into two sets: set 1 and set 2 ii) training the classifier using set 1 and outputting the predicted probability for set 2, and vice versa iii) eventually, after one iteration, we obtained the predicted probability for all samples, and we repeat this for 10 times.

#### C.5 Cross-mice fine-tuning

We first sort 50% of the pretrained dataset, denoted as 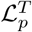. Then we utilize the annotated labels and features from the pretrained dataset together with the new dataset’s annotation to fine-tune the cell classifier. The pretrained classifier is parameterized by *θ*. During the fine-tuning process, the classifier is trained with oversampled concatenation of the pretrained dataset and the newly labeled dataset 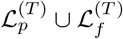 at each iteration *t* to train the new dataset’s cell classifier, parameterized by *θ*′. After training the cell classifier with oversampling techniques, we throw away one sample from the pretrained dataset 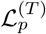 that has the highest classifier uncertainty, defined as 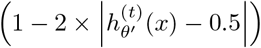. The overall training structure for cross-mice fine-tuning is depicted in Algorithm S2.

##### Algorithm S2 Cross-mice fine-tuning

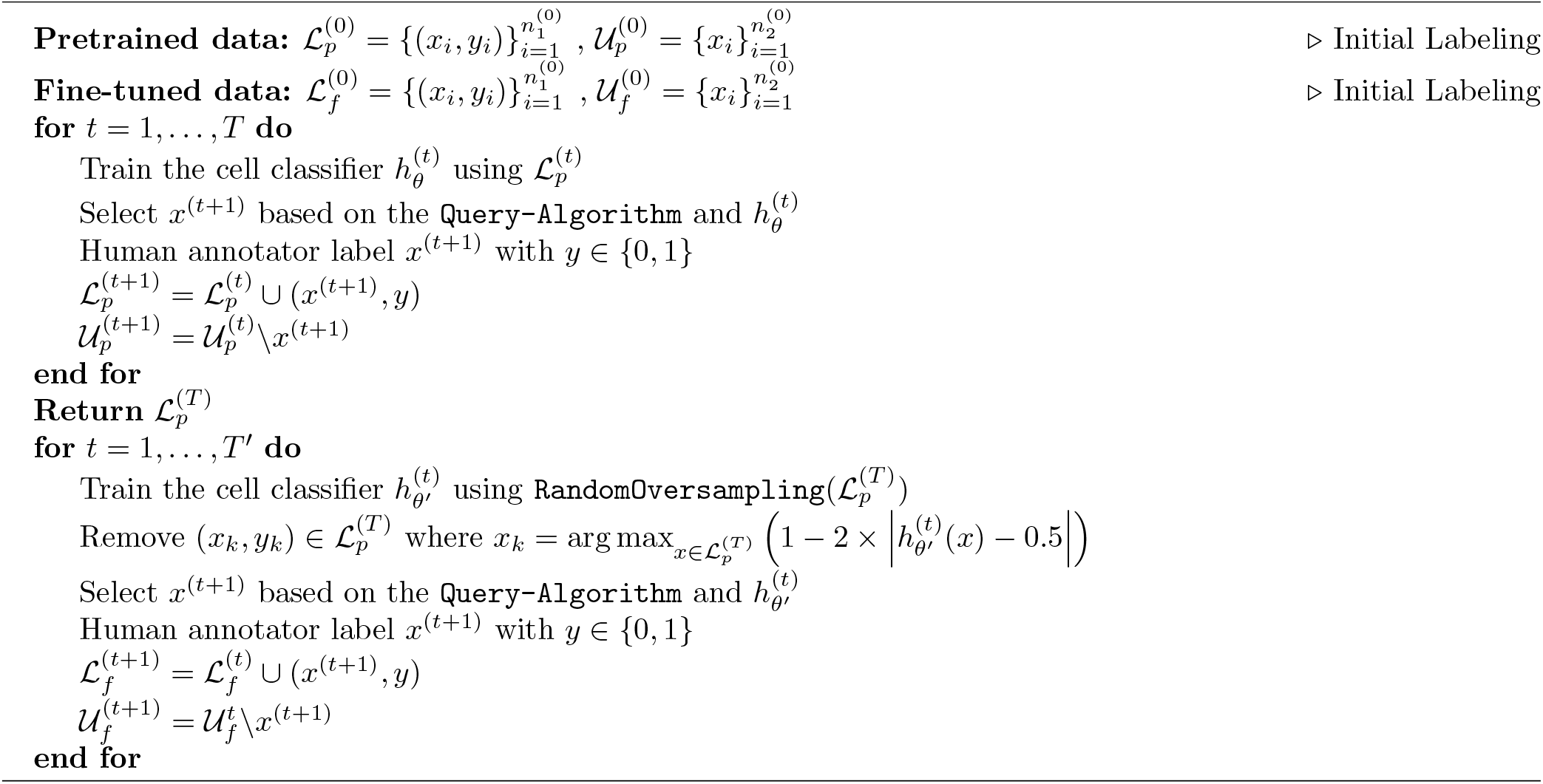

#### C.6 Speed decoding analysis with striatal neurons

To determine whether the quality control process, which aims to remove false-positive cell candidates from the dataset, is essential for subsequent biological analysis, we analyzed calcium imaging data previously acquired from the dorsomedial striatum of freely behaving mice using a head-mounted epifluorescence miniature microscope [56]. We analyzed 9 such movies extracting the Direct Pathway Striatal Projection Neurons (dSPNs, also noted as D1-type neurons) and 10 such movies extracting the Indirect Pathway Striatal Projection Neurons (iSPNs, also noted as D2-type neurons). The cell candidates for each movie are extracted by ICA [64] and have one human annotation for each cell candidate. The speed of each mouse is smoothed with a moving average of window size 30. We then did 1,000 train-test splits (70%-30%) and trained support vector machine regression models to predict the smoothed speeds of the animal using human-annotated cells and ActSort-selected cells, with ActSort trained on 1% of the human annotations. The support vector machine incorporates a *𝓁*1 regularizor with the hyperparameter *λ* selected through a 5-fold cross-validation. We calculated the Pearson correlation coefficient between the predicted speed and the ground truth speed for each dataset using human-rejected candidates, human-selected candidates, and ActSort-selected candidates separately.

### D Resources for reproducibility

Most experiments performed in this work are performed on institution’s clusters, with MATLAB 2022b. Some experiments were performed on a desktop with an Intel(R) Core(TM) i9-10900X CPU with 10 physical cores and an NVIDIA GeForce RTX 4070 Ti GPU, with MATLAB 2021b. If accepted, we will be open-sourcing the software, the benchmark, and the reproduction code to reproduce our results in this paper.

## Figures, Tables, and Captions

**Table S1:** The human annotation benchmark of one-photon (1p) half-hemisphere dataset, 3 mice labeled by 4 human annotators each. The datasets include 28,010 cell candidates and 112,040 annotations. The rows stand for the annotators who were used as ground truth for evaluations, whereas the columns stand for those who are being evaluated. Teal stands for the accuracy of accepting cells that were accepted by the evaluator and red stands for the rejection. Evaluators disagreed on several occasions, leading to average (balanced) accuracy of ∼ 80%. The movie images and the label distributions can be found in Fig S13.

| Ann.<br>Eval. | Guinea pig | Koala | Lion | Panda | Dragon | Cheetah |
| --- | --- | --- | --- | --- | --- | --- |
| Guinea pig | NA | 0.91 0.92 | 0.91 0.88 | 0.97 0.80 | 0.92 0.91 | 0.97 0.69 |
| Koala | 0.99 0.39 | NA | 0.94 0.59 | 0.99 0.39 | NA | 0.98 0.48 |
| Lion | 0.99 0.40 | 0.93 0.63 | NA | 0.98 0.55 | 0.95 0.63 | 0.99 0.32 |
| Panda | 0.99 0.61 | 0.88 0.85 | 0.94 0.76 | NA | 0.92 0.88 | NA |
| Dragon | 0.99 0.45 | NA | 0.93 0.72 | 0.99 0.50 | NA | NA |
| Cheetah | 0.98 0.56 | 0.95 0.70 | 0.87 0.89 | NA | NA | NA |
| Average | 0.99 0.48<br>0.74±0.04 | 0.92 0.78<br>0.85±0.05 | 0.92 0.77<br>0.84±0.05 | 0.98 0.56<br>0.77±0.07 | 0.93 0.81<br>0.87±0.06 | 0.98 0.50<br>0.74±0.07 |

**Table S2:** The human annotation benchmark of one-photon (1p) neocortex dataset with many false positives (2,345 out of 6,691 cell candidates) and 26,764 annotations. The rows stand for the annotators who were used as ground truth for evaluations, whereas the columns stand for those who are being evaluated. Teal stands for the accuracy of acceptance, red stands for the accuracy of rejection. Evaluators disagreed on several occasions, the balanced accuracy between annotators is ∼ 83%. The movie images and the label distribution can be found in Fig S9.

| Ann.<br>Eval. | Guinea pig | Lion | Dragon | Cheetah |
| --- | --- | --- | --- | --- |
| Guinea pig | NA | 0.641 0.972 | 0.831 0.873 | 0.790 0.956 |
| Lion | 0.988 0.432 | NA | 0.976 0.630 | 0.969 0.724 |
| Dragon | 0.959 0.592 | 0.730 0.963 | NA | 0.881 0.906 |
| Cheetah | 0.985 0.561 | 0.783 0.958 | 0.952 0.784 | NA |
| Average | 0.977 0.529<br>0.75±0.09 | 0.718 0.964<br>0.84±0.05 | 0.920 0.762<br>0.84±0.04 | 0.880 0.862<br>0.87±0.03 |

**Table S3:** The human annotation benchmark of two-photon (2p) Ca^2+^ movie with 5,276 cell candidates and 21,104 annotations. The rows stand for the annotators who were used as ground truth for evaluations, whereas the columns stand for those who are being evaluated. Teal stands for the accuracy of acceptance, red stands for the accuracy of rejection. Evaluators disagreed on several occasions, the balanced accuracy between annotators is ∼ 78%. The movie images and the label distribution can be found in Fig S10.

| Ann.<br>Eval. | Cheetah | Panda | Koala | Dragon |
| --- | --- | --- | --- | --- |
| Cheetah | NA | 0.917 0.782 | 0.780 0.850 | 0.888 0.801 |
| Panda | 0.969 0.560 | NA | 0.802 0.783 | 0.921 0.770 |
| Koala | 0.975 0.343 | 0.949 0.441 | NA | 0.924 0.474 |
| Dragon | 0.971 0.492 | 0.953 0.660 | 0.807 0.722 | NA |
| Average | 0.971 0.465<br>0.72±0.1 | 0.940 0.628<br>0.78±0.07 | 0.796 0.785<br>0.80±0.01 | 0.911 0.682<br>0.80±0.06 |

**Table S4:**
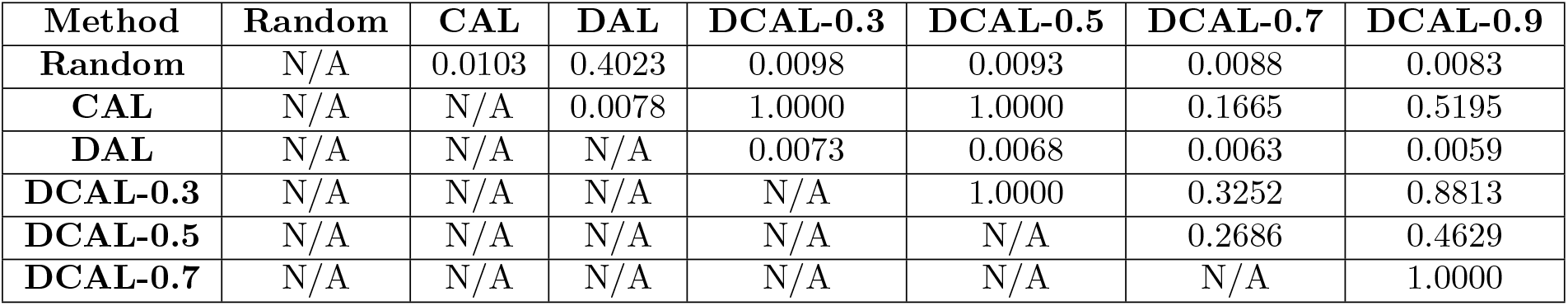
The significance levels for the Wilcoxon signed-rank tests with Bonferroni-Holm corrections performed in Fig. 3B. Each query method had twelve data points (3 datasets and 4 annotators each). For each annotation, we computed the fraction of the boundary samples.

| Method | Random | CAL | DAL | DCAL-0.3 | DCAL-0.5 | DCAL-0.7 | DCAL-0.9 |
| --- | --- | --- | --- | --- | --- | --- | --- |
| Random | N/A | 0.0103 | 0.4023 | 0.0098 | 0.0093 | 0.0088 | 0.0083 |
| CAL | N/A | N/A | 0.0078 | 1.0000 | 1.0000 | 0.1665 | 0.5195 |
| DAL | N/A | N/A | N/A | 0.0073 | 0.0068 | 0.0063 | 0.0059 |
| DCAL-0.3 | N/A | N/A | N/A | N/A | 1.0000 | 0.3252 | 0.8813 |
| DCAL-0.5 | N/A | N/A | N/A | N/A | N/A | 0.2686 | 0.4629 |
| DCAL-0.7 | N/A | N/A | N/A | N/A | N/A | N/A | 1.0000 |

**Table S5:** The significance levels for the Wilcoxon signed-rank tests with Bonferroni-Holm corrections performed in Fig. 3D. Each query method had twelve data points (3 datasets and 4 annotators each). For each annotation, we computed the average cosine distances between the boundary samples. To facilitate a fair comparison, for each query algorithm, we randomly subsampled the number of boundary samples to match the query algorithm with the lowest number of boundary samples.

| Method | Random | CAL | DAL | DCAL-0.3 | DCAL-0.5 | DCAL-0.7 | DCAL-0.9 |
| --- | --- | --- | --- | --- | --- | --- | --- |
| Random | N/A | 1.0000 | 0.0103 | 1.0000 | 0.8145 | 0.6172 | 0.6460 |
| CAL | N/A | N/A | 0.0098 | 0.4702 | 0.0093 | 0.0146 | 0.0479 |
| DAL | N/A | N/A | N/A | 0.0088 | 0.0083 | 0.0444 | 0.0078 |
| DCAL-0.3 | N/A | N/A | N/A | N/A | 0.7568 | 0.0820 | 0.2310 |
| DCAL-0.5 | N/A | N/A | N/A | N/A | N/A | 0.3418 | 0.7764 |
| DCAL-0.7 | N/A | N/A | N/A | N/A | N/A | N/A | 0.8501 |

**Table S6:**
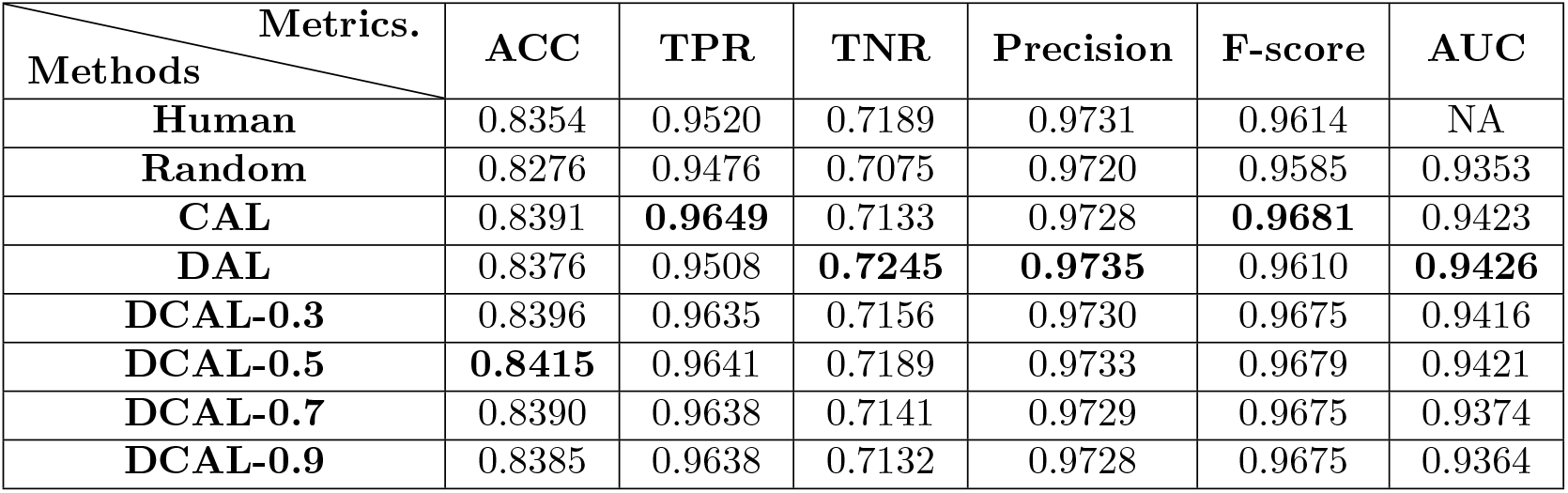
ActSort performance after sorting 5% of the half-hemisphere Ca^2+^ datasets (across 3 mice). Rows, evaluation metrics, where ACC is balanced accuracy, TPR is true positive rate, TNR is true negative rate, and AUC is the area under the curve. Columns, active learning method, where the fraction next to DCAL refers to the corresponding weight. Each entry shows the average metric value for sorting up to 5% across 12 annotators over three datasets. The entire sorting performance as a function of the percentage of sorted cells up to 50% is illustrated in Figs. 4**B-D** and S13**B**.

| Methods \ Metrics. | ACC | TPR | TNR | Precision | F-score | AUC |
| --- | --- | --- | --- | --- | --- | --- |
| Human | 0.8354 | 0.9520 | 0.7189 | 0.9731 | 0.9614 | NA |
| Random | 0.8276 | 0.9476 | 0.7075 | 0.9720 | 0.9585 | 0.9353 |
| CAL | 0.8391 | **0.9649** | 0.7133 | 0.9728 | **0.9681** | 0.9423 |
| DAL | 0.8376 | 0.9508 | **0.7245** | **0.9735** | 0.9610 | **0.9426** |
| DCAL-0.3 | 0.8396 | 0.9635 | 0.7156 | 0.9730 | 0.9675 | 0.9416 |
| DCAL-0.5 | **0.8415** | 0.9641 | 0.7189 | 0.9733 | 0.9679 | 0.9421 |
| DCAL-0.7 | 0.8390 | 0.9638 | 0.7141 | 0.9729 | 0.9675 | 0.9374 |
| DCAL-0.9 | 0.8385 | 0.9638 | 0.7132 | 0.9728 | 0.9675 | 0.9364 |

**Table S7:** ActSort performance on 2% of the batch sorted cell candidates. Rows, evaluation metrics, where ACC is balanced accuracy, TPR is true positive rate, TNR is true negative rate, and AUC is the area under the curve. Columns, active learning method, where the fraction next to DCAL refers to the corresponding weight. Each entry shows the average metric value for sorting up to 2% across 64 augmented annotators. The entire sorting performance as a function of the percentage of sorted cells up to 5% is illustrated in Figs. 4**E-G** and S13**C**.

| Metrics.<br>Methods | ACC | TPR | TNR | Precision | F-score | AUC |
| --- | --- | --- | --- | --- | --- | --- |
| **Human** | 0.8354 | 0.9520 | 0.7189 | 0.9731 | 0.9614 | NA |
| **Random** | 0.8349 | 0.9545 | 0.7153 | 0.9733 | 0.9634 | 0.9391 |
| **CAL** | 0.8398 | 0.9715 | 0.7080 | 0.9730 | 0.9720 | 0.9434 |
| **DAL** | 0.8373 | 0.9601 | 0.7144 | 0.9733 | 0.9663 | **0.9453** |
| **DCAL-0.3** | 0.8431 | **0.9716** | 0.7146 | 0.9736 | 0.9723 | 0.9436 |
| **DCAL-0.5** | 0.8444 | 0.9713 | 0.7175 | 0.9738 | 0.9723 | 0.9439 |
| **DCAL-0.7** | **0.8465** | 0.9708 | **0.7222** | **0.9742** | **0.9723** | 0.9444 |
| **DCAL-0.9** | 0.8455 | 0.9703 | 0.7208 | 0.9741 | 0.9719 | 0.9433 |

**Table S8:** ActSort performance on fine-tuning experiment with 1% annotation. Rows, evaluation metrics, where ACC is balanced accuracy, TPR is true positive rate, TNR is true negative rate, and AUC is the area under the curve. Columns, active learning method, where the fraction next to DCAL refers to the corresponding weight. Each entry shows the average metric value for sorting up to 1% across 96 augmented annotators over 6 dataset pairs. The experimental details are described in Fig. 5 and the entire sorting performance as a function of the percentage of sorted cells up to 20% are illustrated in Fig. 5 **B-D** and Fig. S13**D**.

| Metrics.<br>Methods | ACC | TPR | TNR | Precision | F-score | AUC |
| --- | --- | --- | --- | --- | --- | --- |
| **Human** | 0.8354 | 0.9520 | 0.7189 | 0.9731 | 0.9614 | NA |
| **Random** | 0.8112 | 0.9363 | 0.6860 | 0.9702 | 0.9513 | 0.9208 |
| **CAL** | 0.8261 | **0.9551** | 0.6970 | 0.9714 | **0.9621** | 0.9131 |
| **DAL** | 0.8300 | 0.9346 | **0.7254** | 0.9735 | 0.9521 | **0.9335** |
| **DCAL-0.3** | 0.8323 | 0.9515 | 0.7130 | 0.9727 | 0.9608 | 0.9170 |
| **DCAL-0.5** | 0.8337 | 0.9509 | 0.7165 | 0.9730 | 0.9606 | 0.9186 |
| **DCAL-0.7** | 0.8346 | 0.9509 | 0.7183 | 0.9731 | 0.9607 | 0.9194 |
| **DCAL-0.9** | **0.8364** | 0.9503 | 0.7225 | **0.9735** | 0.9605 | 0.9211 |

**Figure S1:**
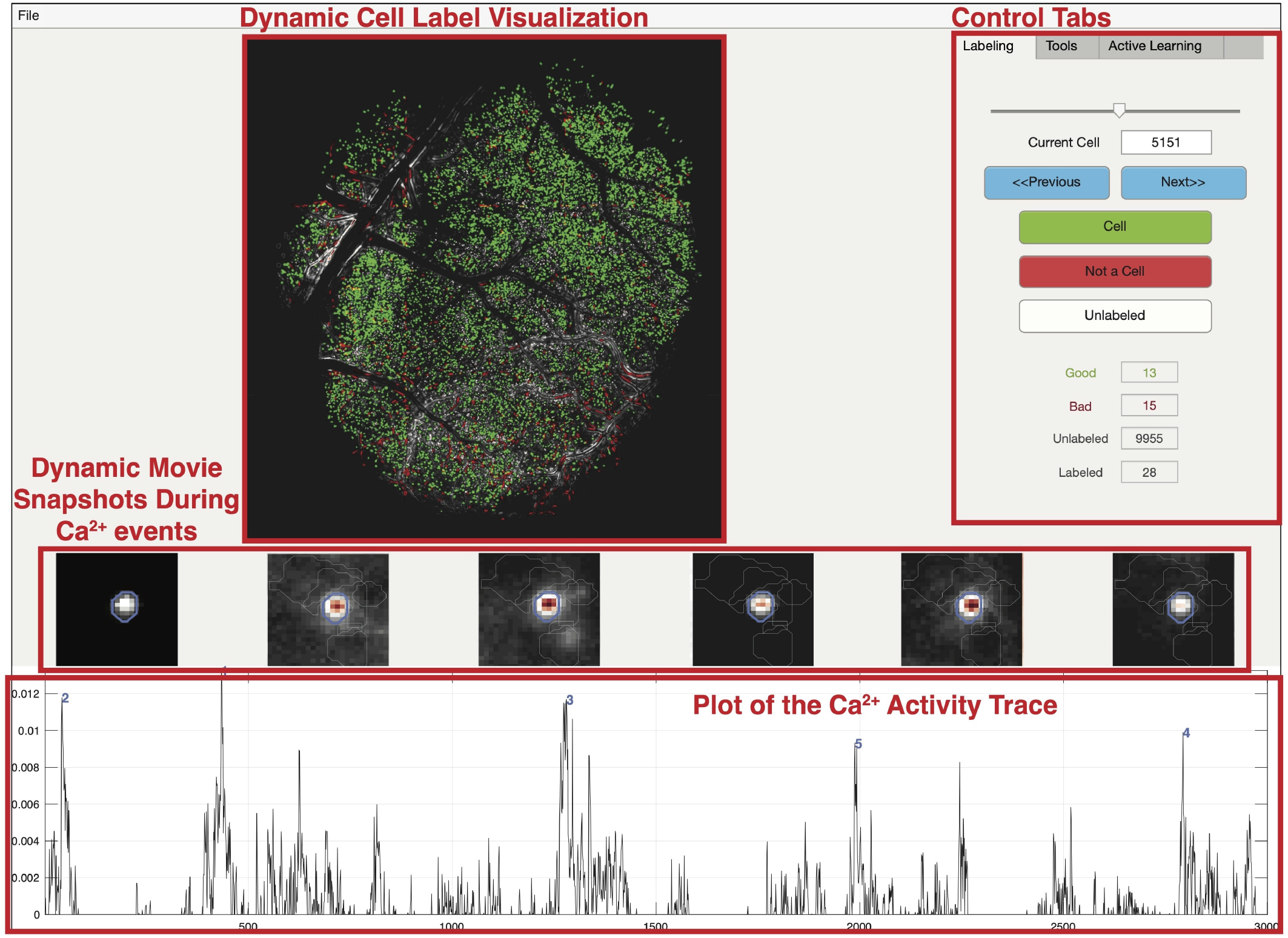
ActSort has a user-friendly cell sorting software for experimental neuroscientists. The user interface of ActSort offers options to watch movie snapshots, study Ca^2+^ activity traces, zoom into the maximum projection image of the movie, and has several other controls detailed in Appendix A. ActSort is available on MATLAB apps and can be used on personal laptop thanks to the data compression module.

**Figure S2:**
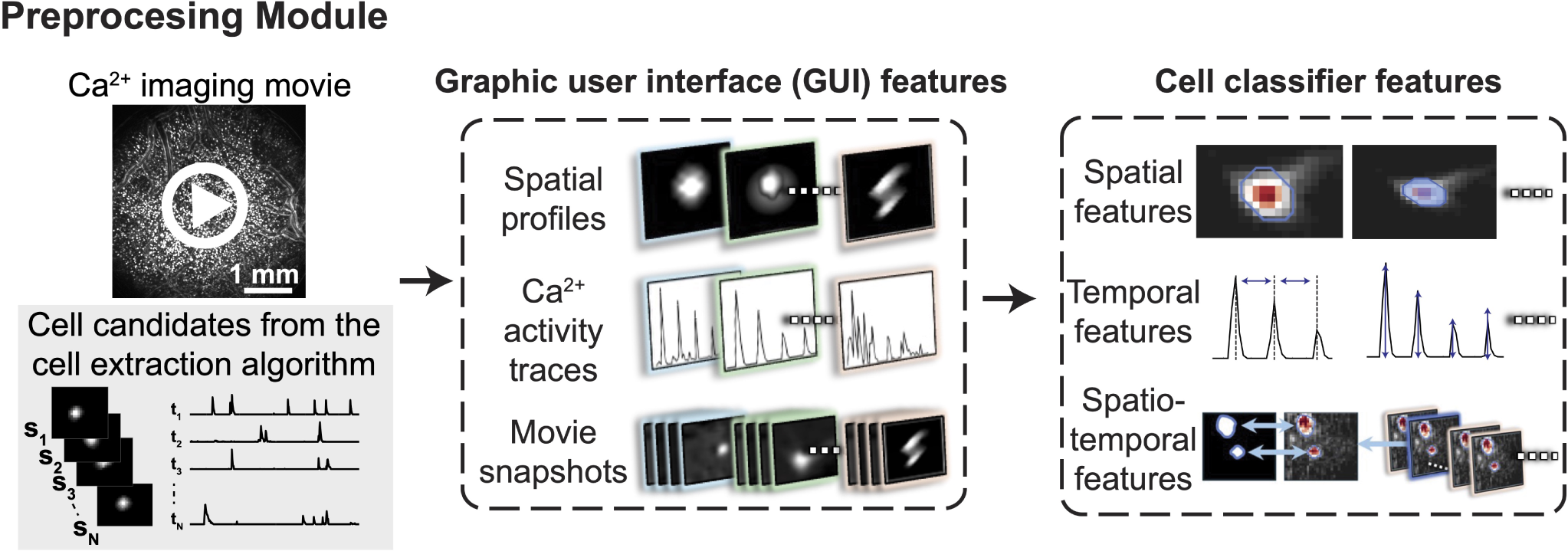
A visualization of ActSort’s preprocessing module. In response to the challenge of managing rapidly increasing file sizes in Ca^2+^ imaging movies and cell extraction results —often reaching TBs— the preprocessing module is crucial for enabling memory efficient real-time processing. This module efficiently compresses the input data to just a few GBs, facilitating easy data transfer and use on personal laptops. *Left*. The inputs to the preprocessing module include the Ca^2+^ imaging movie and the cell extraction results. *Middle*. The processed data are transformed into GUI-compatible features such as spatial profiles, Ca^2+^ activity traces, and Ca^2+^ movie snapshots. These features are then utilized by the GUI module to support human annotation decisions. *Right*. The input data are converted into features, quality metrics that quantify the properties of individual cells (**Method**; Appendix A.1). These features are essential for training the classifiers later in the pipeline.

**Figure S3:**
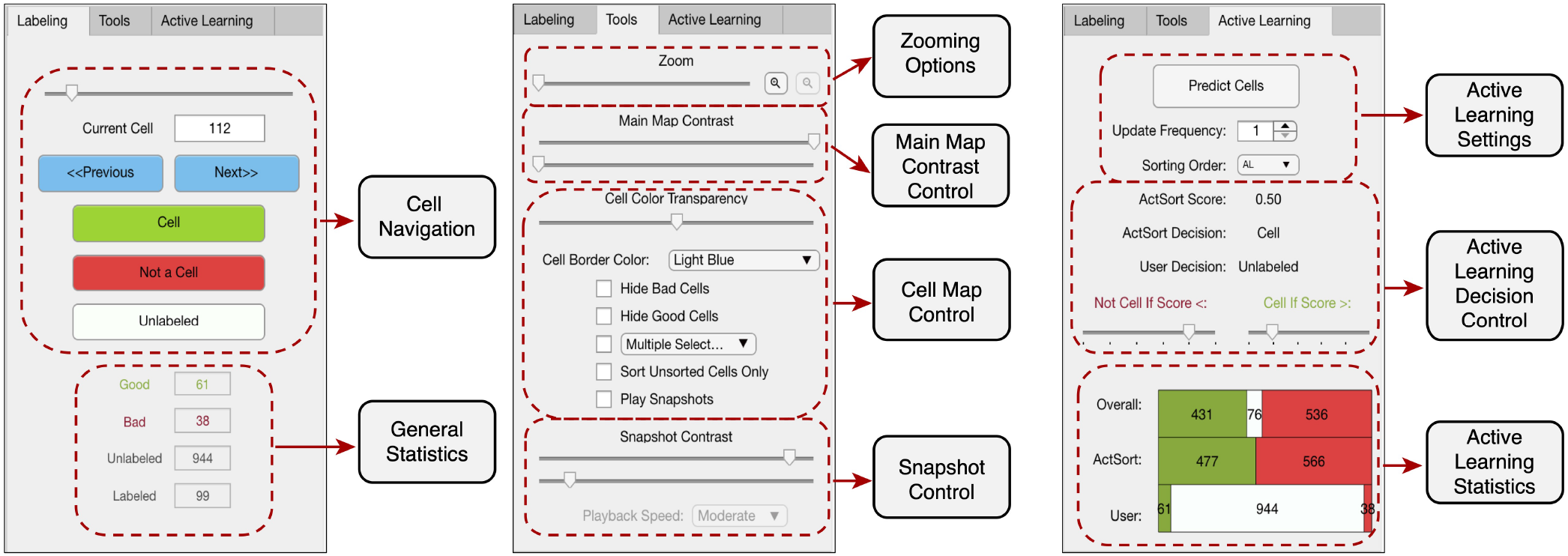
The control tabs within the ActSort software. A visualization highlighting the broad functionalities of the ActSort software. *Left*. In the labeling tab, the user can navigate between cells using a slider and make decisions by clicking on buttons. These can also be achieved by keyboard shortcuts and by clicking on the cells from the cell map (Fig. S1). *Middle*. The tools tab allows the user to zoom into the map, which can be performed via sliders or by clicking with the mouse. The user also has the option to adjust the map/snapshot contrasts, perform multiple selections, and many other utilies discussed in Appendix A. *Right*. The active learning tab allows the user to monitor the progress of the active learning algorithm. The user has the option to let the active learning make the decisions for the unsorted cells, which can be previewed on the cell map. Finally, the training frequency of the decoders can be decreased for exceptionally large datasets to allow faster sorting. This feature was designed mainly to allow scalability and further development, *i*.*e*., for future active learning algorithms that may need more time to infer the next query.

**Figure S4:**
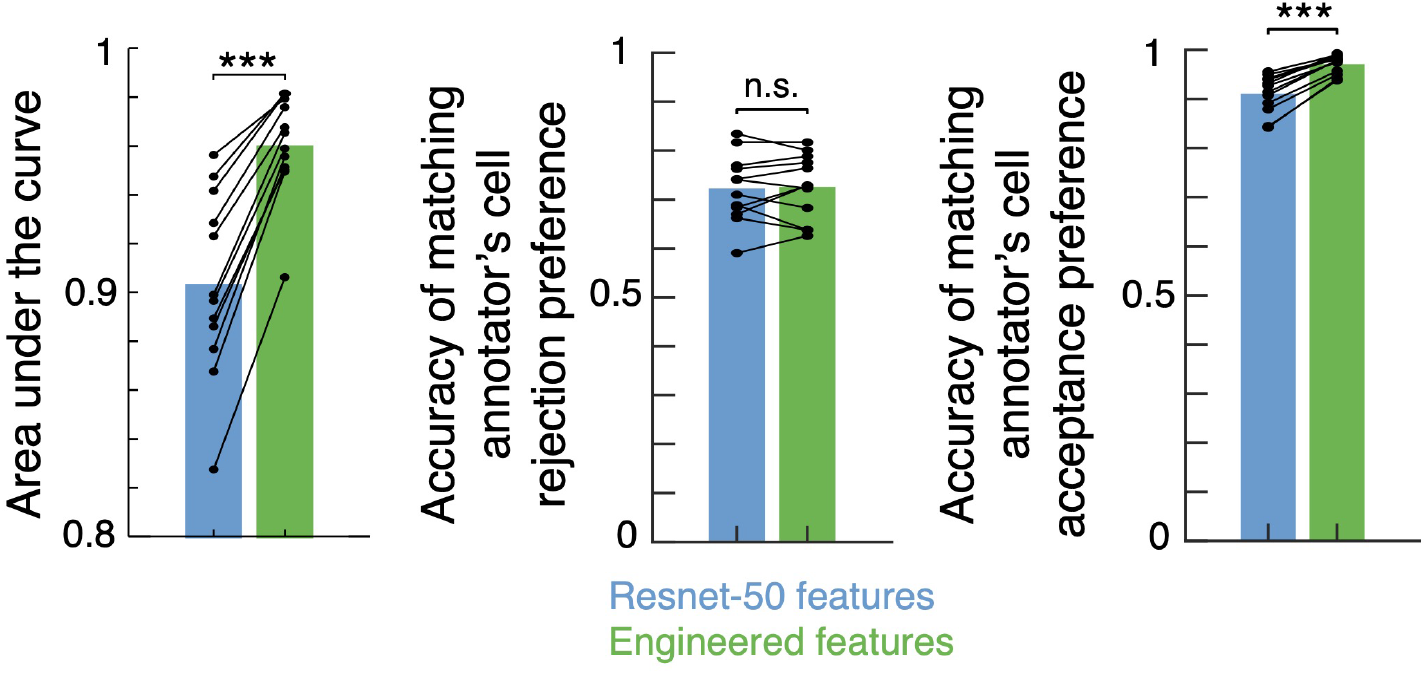
Features derived from Resnet-50 failed to match the performance of expert-engineered features. To test whether pre-trained deep networks can be used as cell classifiers, we once again used the half-hemisphere dataset. We ran cropped Ca^2+^ imaging movie snapshots, averaged across frames at the time of cells’ Ca^2+^ events, and cells’ extracted spatial footprints on a pre-trained Resnet-50 model. We extracted the resulting 2,000 features from the penultimate layer, which we term as “ResNet-50 features.” *Left*. Similar to Fig. 2**A**, we trained cell classifiers (**Method**; Appendix C.1) on the engineered and ResNet-50 features. Expert-engineered features outperformed ResNet-50 features, leading to higher area under the receiver operating curve. *Middle*. The true negative rates obtained from the engineered or ResNet-50 features were mostly identical. *Right*. The expert-engineered features led to higher true positive rates, whereas ResNet-50 features rejected 10% of true cells, far below the human performance. Overall, our engineered features outperformed the features extracted from the Resnet-50 model. Each dot corresponds to a single annotation instance. All tests are two-sided Wilcoxon signed-rank tests (^***^*p* < 10^−3, **^*p* < 10^−2, *^*p* < 0.05).

**Figure S5:**
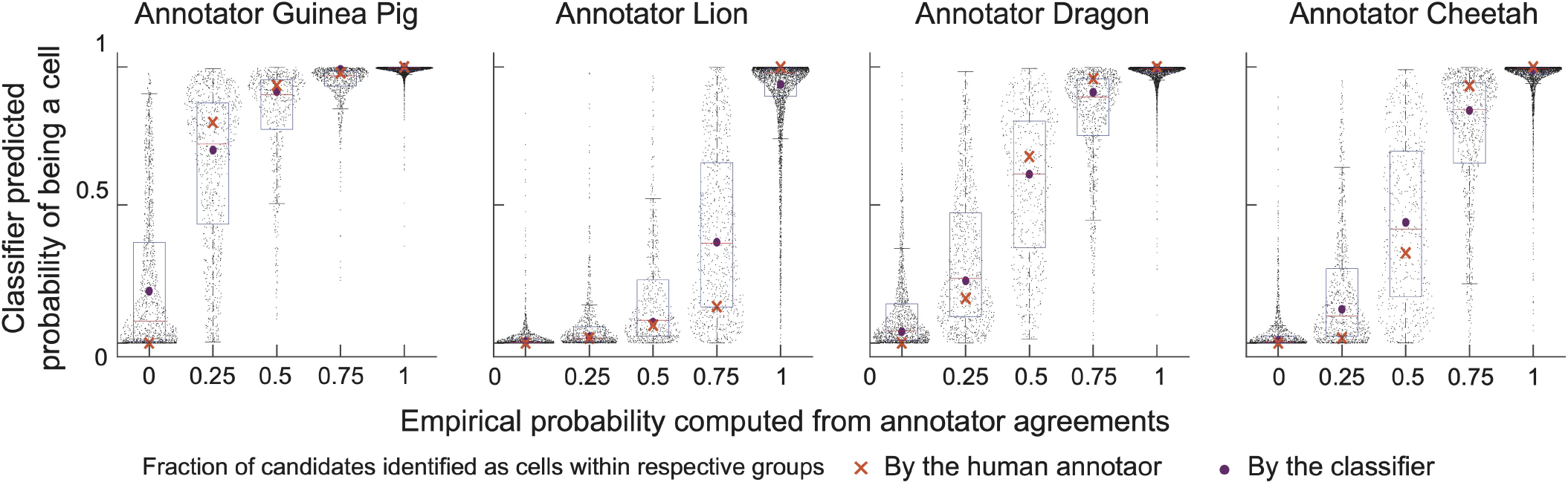
Cell classifiers can mitigate human annotator bias. We trained the cell classifier on the neocortex datasets (**Method**; Appendix C.1) and predicted the test probabilities for each cell candidate. The empirical probability for each cell is computed from annotator agreements (as discrete values: 0, 0.25, 0.5, 0.75, 1). Each plot corresponds to an annotator. The performance for each annotator can be found in Table S2. Each dot represents a single cell from the neocortical dataset [44]. Red lines, median; purple dots, mean of the corresponding distribution; red cross, mean of the corresponding human annotations’ distribution. The classifier shows a better-calibrated prediction, while the human annotators (especially first two researchers) display strong biases.

**Figure S6:**
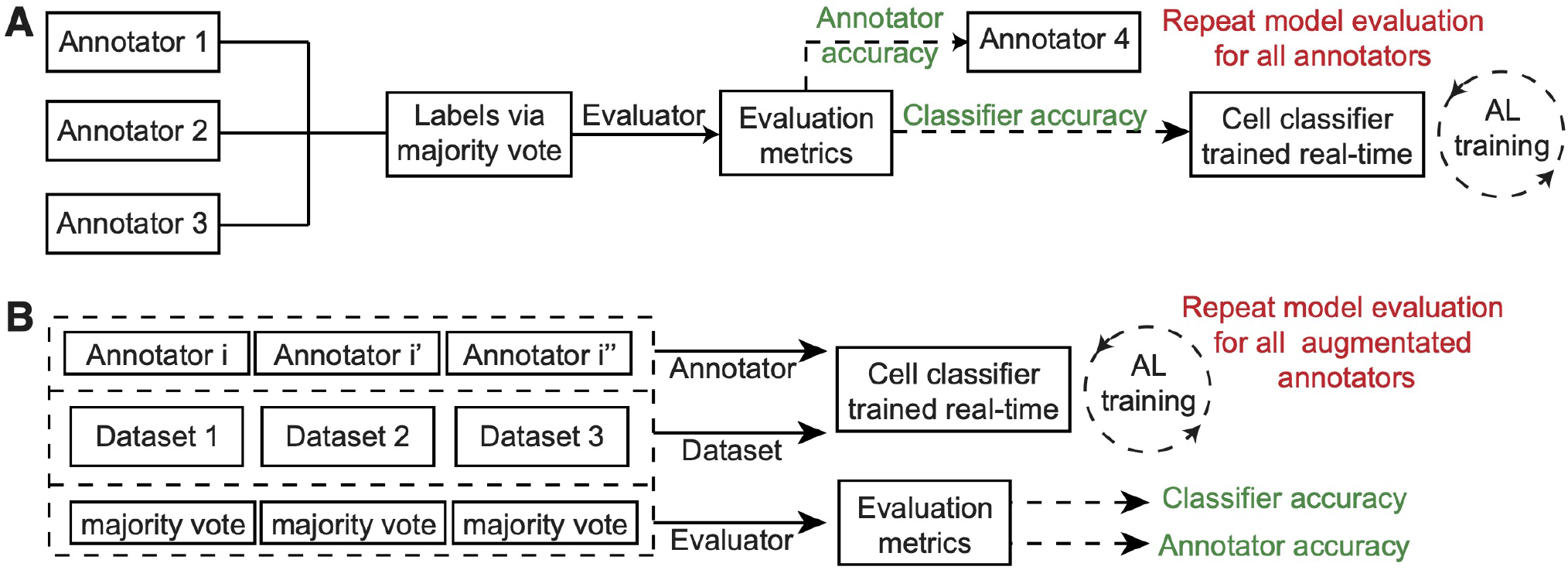
A description of the evaluation methods we used in the main text for benchmarking. **A** Our workflow for testing online classifier’ accuracies when trained with the labels from a distinct annotator. The majority votes from the rest of the annotators are used in place of the ground truth, which we call “evaluators.” This workflow is used in all the experiments in Sections 2.8 and 2.9. In the cross-mice experiment, Section 2.9, the creation of annotator and evaluator is repeated twice for both the pre-training stage and the fine-tuning stage. **B** When augmenting new annotators for experiments involving multiple imaging datasets, we first computed the evaluators through the majority vote for each annotator and concatenated both to obtain augmented annotators and evaluators. This way of augmentation is used for Figs. 4**E-G** and S13**C**.

**Figure S7:**
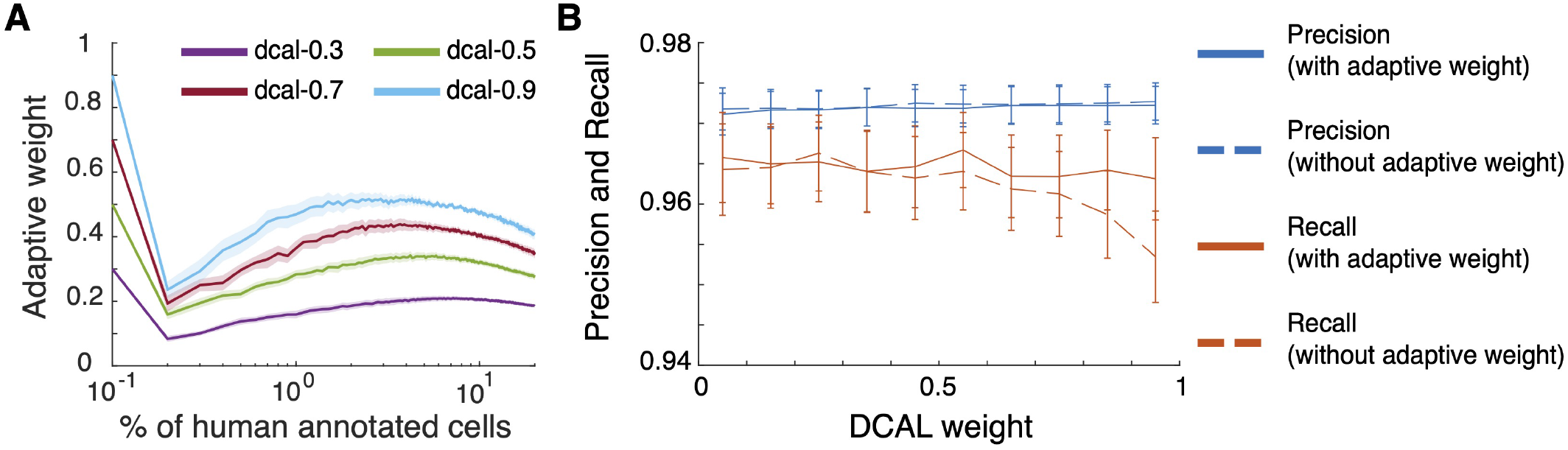
Adaptive estimation of the model parameter leads to improved true positive rates. To study the role of the adaptive weight estimation in DCAL, we visualized the evolution of the weight parameter throughout the sorting process of the half-hemisphere dataset and quantified the effectiveness of the adaptive weight strategy. **A** The adaptive weight value was recorded as we ran ActSort on the half-hemisphere dataset, sorting up to 20%. The sudden drop in the weight values when only a few samples are sorted is due to insufficient training data, leading to class imbalance. Fortunately, the adaptive estimation decreases the sensitivity to the initial choices of *w*. Solid lines: means. Shaded areas: s.e.m. over 12 annotators. **B** We computed the precision and recall values after running ActSort with DCAL (with and without adaptive estimation) on the half-hemisphere dataset, sorting up to 3% cell candidates. The weights were initially swept within the interval [0.02, 0.98], and later binned with a window size of 0.1 due to high variability and low number of annotations. Solid lines: precision (blue) and recall (also known as true positive rate, red) values with adaptive weight estimation. Dashed lines: The same, but without adaptive estimation. Without adaptive weight estimation, DCAL became suboptimal with increased weight values, leading to decreased true positive rates.

**Figure S8:**
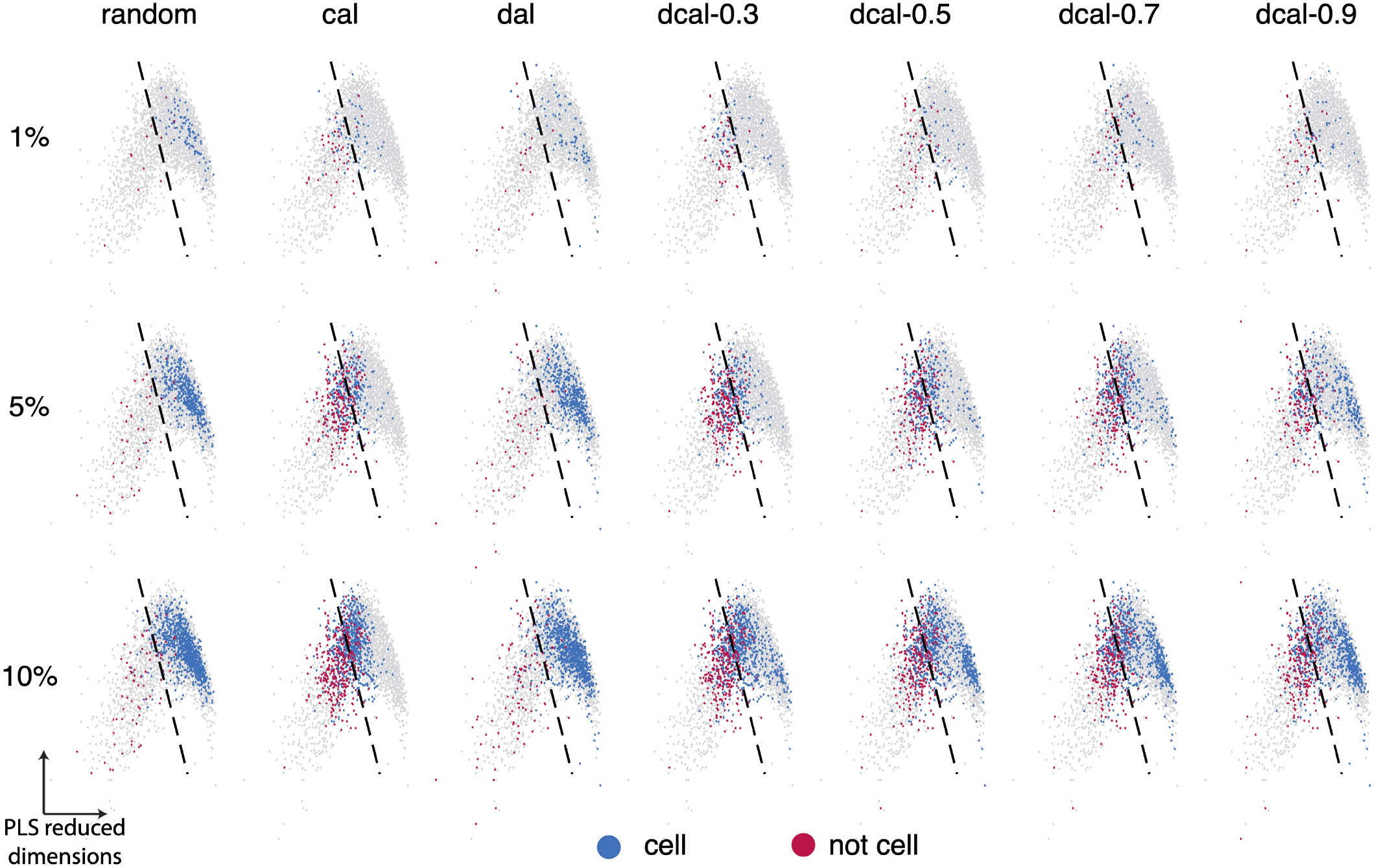
The distribution of selected samples from various query algorithms. This is an extended visualization for Fig. 3**A**, but for 1%, 5%, and 10% annotated datasets.

**Figure S9:**
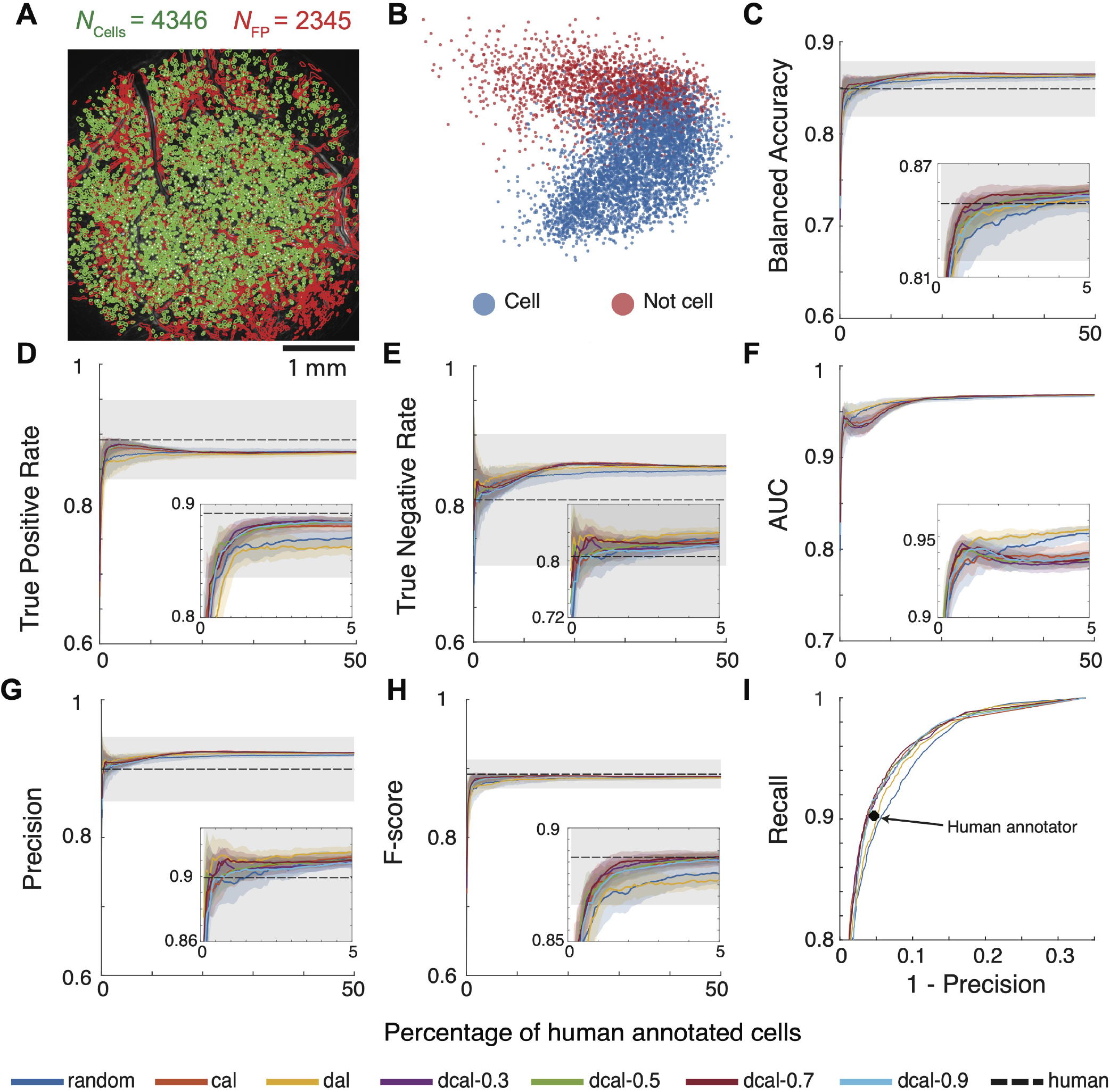
ActSort improves cell sorting quality in movies with many false positives. To assess the generalization ability of ActSort in one-photon Ca^2+^ imaging movies with many false positives, we ran the active learning experiments on the neocortex dataset (**Method**; Appendix B). **A** Example cell map of the neocortical brain regions [44]. Green lines indicate cell boundaries, while the red line indicates false positives. **B** Supervised partial least squares (PLS) plots for all samples, similar to Fig. 3**A**, illustrating the separability of true and false positives by the engineered features. **C-H** The results of the active learning-based training on individual annotations. **I** Example precision-recall (PR) curve for the best annotator’s annotation instance. The best annotator is selected as the one who has the highest average accuracy from Table S2. Solid lines: means. Shaded areas: s.e.m. over four annotators. Dashed lines and gray areas: mean and s.e.m. from four human annotations.

**Figure S10:**
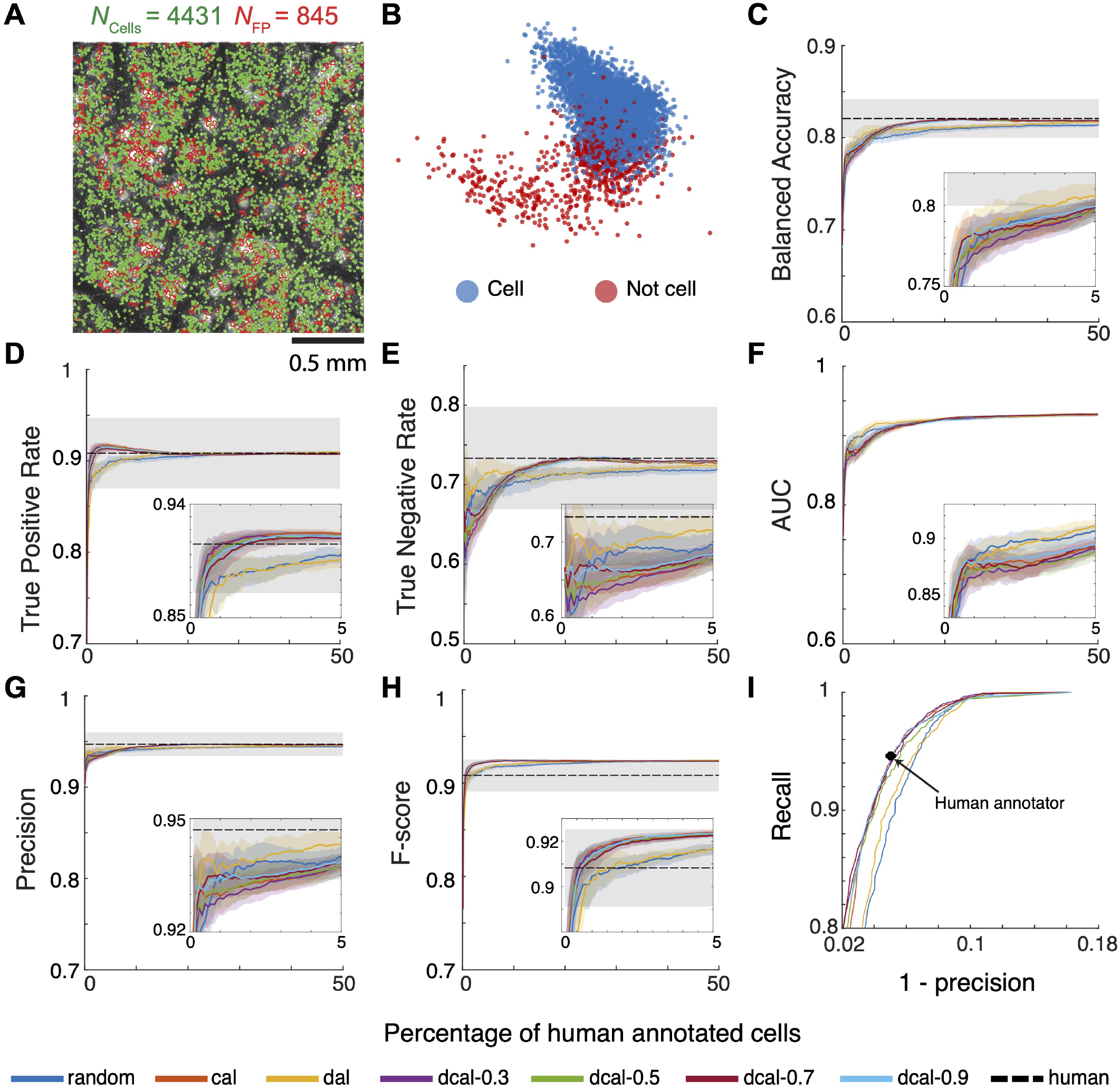
ActSort can process two-photon Ca^2+^ imaging movies with residual motion. With ActSort, we processed an example two-photon Ca^2+^ imaging movie recording layer 2/3 pyramidal cells in primary visual cortex (**Method**; Appendix B). The movie had uncorrected residual motion by design, which led to several false positives in the form of duplicates. **B-I** Same as Fig. S9.

**Figure S11:**
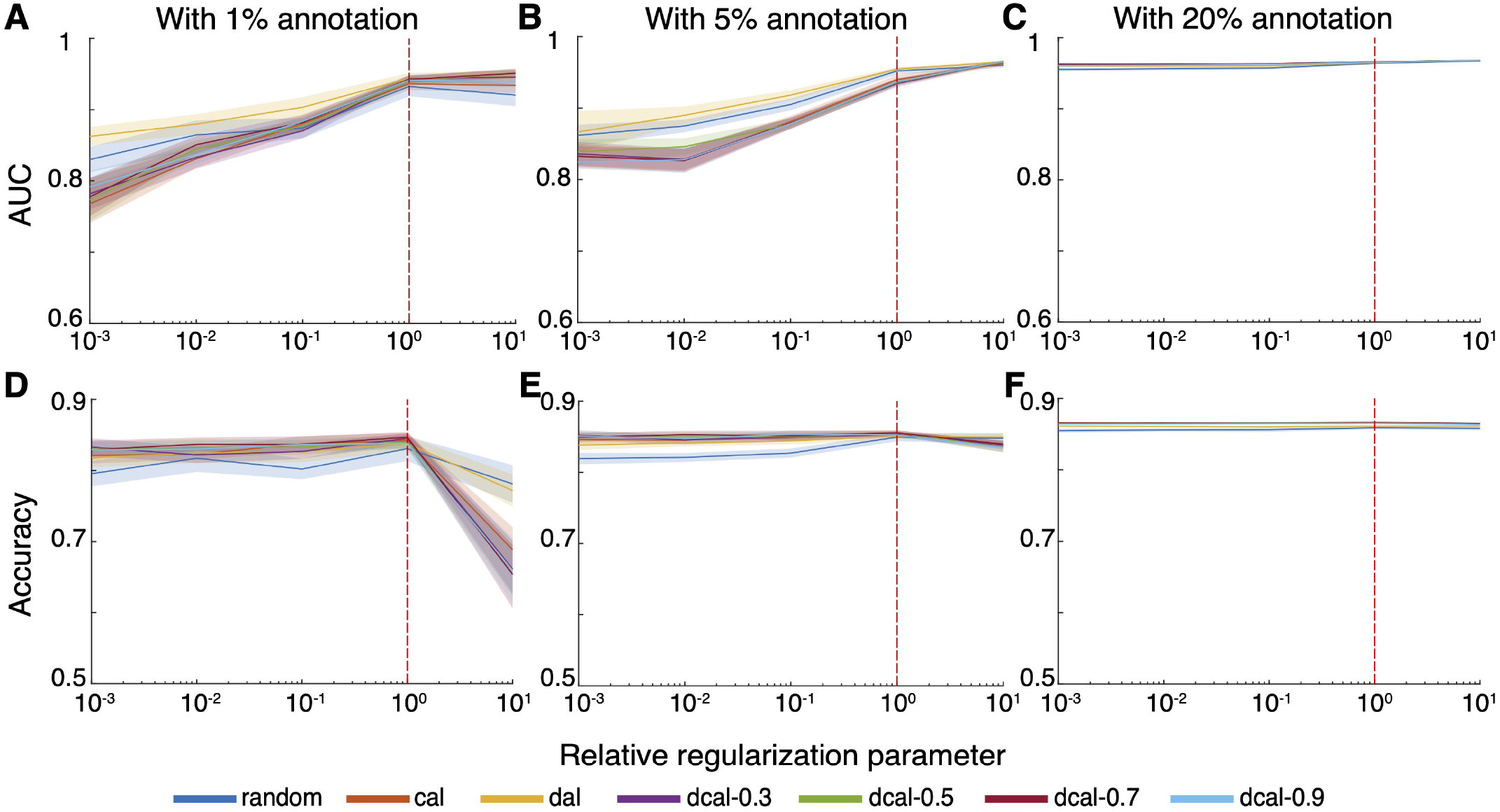
The insensitivity to the regularization strength allows rapid training without cross-validation. Our emphasis on real-time training necessitated using a pre-defined regularization strength, which was taken to be *λ*_default_ = 1*/*(Number of Samples). To validate this, we reanalyzed the neocortex dataset with varying levels of regularization parameters for the cell classifiers. The first row illustrates the area under the receiver operating curve (AUC) for the cell classifier’s predictions at different stages of annotation—1%, 5%, and 20% of cell candidates—with *λ*_relative_ = 1 corresponds to the default value for the problem of interest. The second row shows the balanced accuracy of the classifier’s predictions at these same annotation stages. Solid lines: means. Shaded areas: s.e.m. over four annotators. Red dashed lines: performance metrics at the default regularization value.

**Figure S12:**
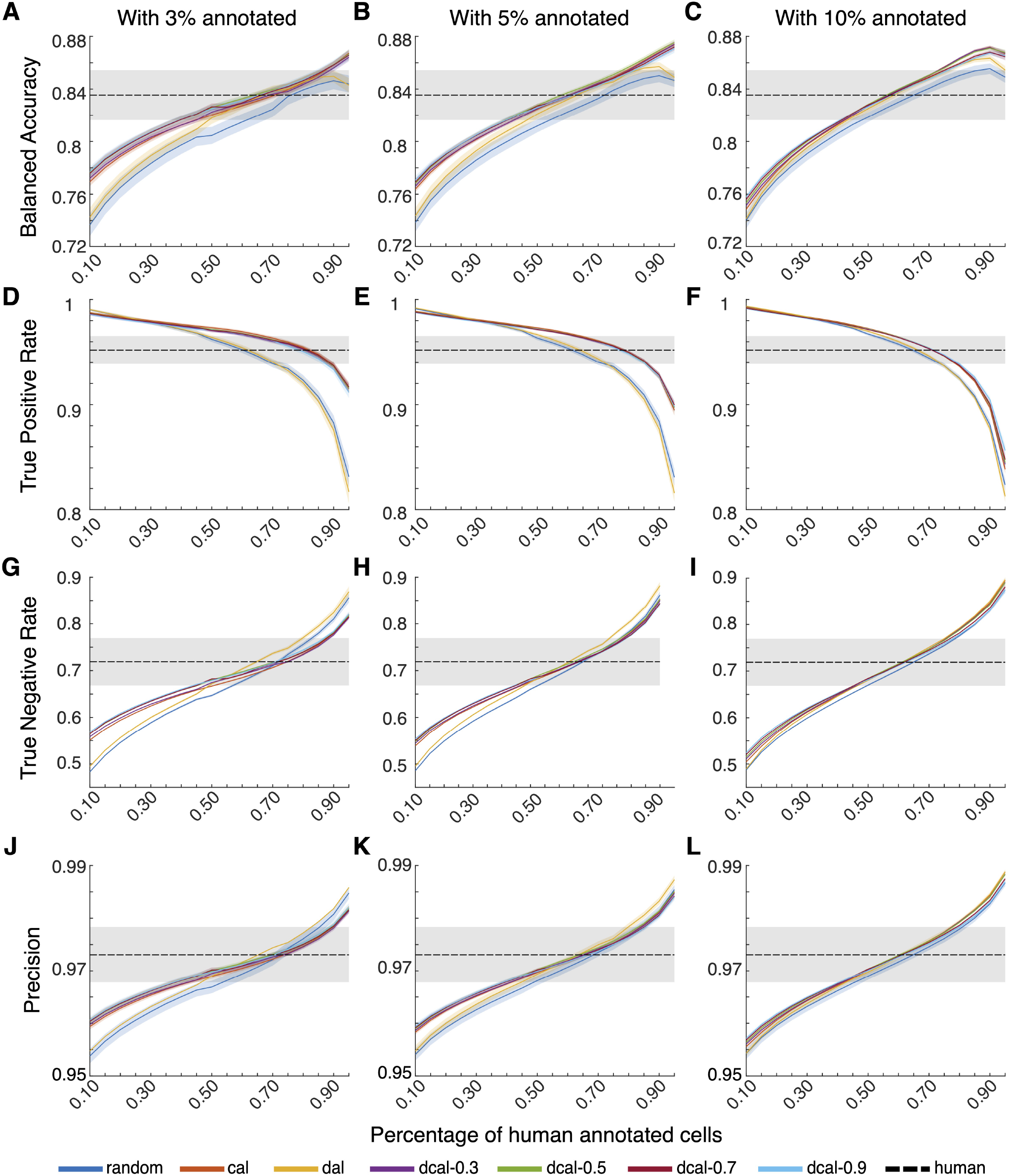
ActSort performance on various classifier thresholds. A sampled collected from the decision boundary should be equally likely to be true or false positive. In contrast, we found that human annotators tend to reject cell candidates close to the decision boundary, diminishing the efficacy of the default 0.5 threshold for the cell classifier. The need for real-time training compelled us to establish a predefined classifier threshold. We conducted additional active learning experiments on the half-hemisphere dataset and assessed the classifier’s balanced accuracy, true positive rate, true negative rate, and precision at different annotation stages—1%, 5%, and 10%-under different classifier threshold, ranging from 0.1 to 0.95. In almost all cases, a threshold between 0.6 and 0.7 aligned well with human annotation preferences. Solid lines: means. Shaded areas: s.e.m. over 12 annotators. Dashed lines and gray areas:, means and s.e.m. from human annotations.

**Figure S13:**
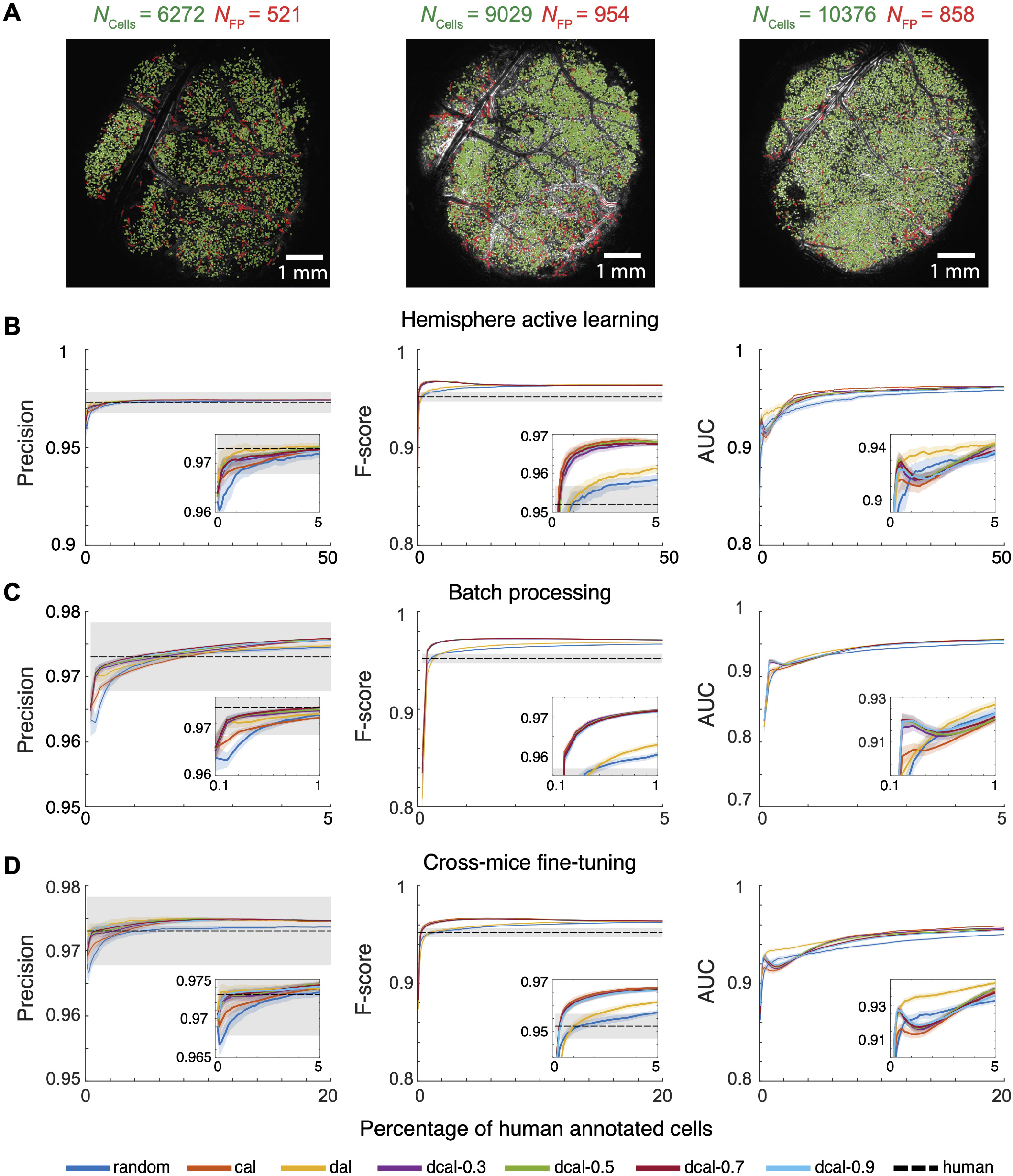
ActSort performance on various types of experiments outperform human annotators. **A** Example cell map from 7mm window half-hemisphere across three mice (**Method**; Appendix B). Green lines, cell boundaries; red lines, false-positives. **B-D** The remaining evaluation metrics for (**B**) Fig. 4**B-D**, (**C**) Fig. 4**E-G**, and (**D**) Fig. 5, including recall, precision, F-score, and area under the receiver operating characteristic curve.

## Footnotes

1 ^1^For the purpose of this work and real-time processing in the software, we used a fixed default value for the regularization, which we validated in Fig. S11.

## Notes

### Competing Interest Statement

The authors have declared no competing interest.

